# Enzyme activation by urea reveals the interplay between conformational dynamics and substrate binding: a single-molecule FRET study

**DOI:** 10.1101/2024.09.01.610662

**Authors:** David Scheerer, Dorit Levy, Remi Casier, Inbal Riven, Hisham Mazal, Gilad Haran

## Abstract

Proteins often harness extensive motions of domains and subunits to promote their function. Deciphering how these movements impact activity is key for understanding life’s molecular machinery. The enzyme adenylate kinase is an intriguing example for this relationship; it ensures efficient catalysis by large-scale domain motions that lead to the enclosure of the bound substrates ATP and AMP. At high concentrations, AMP also operates as an allosteric inhibitor of the protein. Surprisingly, the enzyme is activated by urea, a compound commonly acting as a denaturant. Combining single-molecule FRET spectroscopy and enzymatic activity studies, we find that urea interferes with two key mechanisms that contribute to enzyme efficacy. First, urea promotes the open conformation of the enzyme, aiding the proper positioning of the substrates. Second, urea decreases AMP affinity, paradoxically facilitating a more efficient progression towards the catalytically active complex. These results signify the important interplay between conformational dynamics and chemical steps, including binding, in the activity of enzymes. State-of-the-art tools, such as single-molecule fluorescence spectroscopy, offer new insights into how enzymes balance different conformations to regulate activity.

## Introduction

Enzymes accelerate vital chemical reactions by many orders of magnitude. To promote and regulate their activity, some enzymes have evolved to harness large-scale motions of domains and subunits. Understanding the role of conformational dynamics is crucial for deciphering the functionality of such proteins, as identified in multiple experimental and theoretical studies.^1–4^ One paradigmatic example of a strong relation between conformational dynamics and activity is adenylate kinase (AK),^5–8^ which plays a key role in maintaining cell ATP levels by catalyzing the reaction ATP + AMP ⇄ ADP + ADP.^9, 10^ X-ray crystallographic studies^11–13^ showed that the three-domain enzyme undergoes a significant conformational rearrangement upon substrate binding. The LID domain and nucleotide monophosphate (NMP)-binding domain of the protein close in over the large CORE domain, forming the active center and excluding solvent molecules that might interfere with the chemical reaction. AK’s domain-closure dynamics have been studied using NMR spectroscopy,^6–8, 14^ bulk^5^ and single-molecule fluorescence experiments,^5, 6, 15–17^ as well as multiple molecular dynamics simulations.^18–24^ The dynamics can be described in terms of two states, open and closed. While the open conformation dominates in the absence of substrates, the population of the closed state increases upon substrate binding. Interestingly, a quantitative analysis of the interconversion rates between the two states, based on single-molecule FRET (smFRET) experiments, demonstrated that AK’s conformational dynamics are significantly faster than its turnover,^16, 17^ a finding supported by several computational studies.^20, 25–28^ Indeed, in the presence of substrates, domain closing and opening are found to complete in just a few tens of microseconds. It has been proposed that these fast domain movements might assist the enzyme in orienting substrates for catalysis.^25^ A recent model that combines conformational dynamics and biochemical steps explains the complex activity of AK, particularly the inhibition by its own substrate, AMP, at high concentrations.^17^

An intriguing phenomenon of AK is the enhancement of activity by the denaturant urea at concentrations well below protein denaturation.^29–31^ Originally, it was suggested that the activation is based on increased conformational flexibility at the active site.^29^ In contrast, nuclear magnetic resonance (NMR) experiments suggested that the activation might be due to a redistribution of structural states.^30^ These studies primarily focused on individual substrate concentrations and addressed the role of conformational changes only indirectly. In the current work, we provide a complete study of the effects of the denaturant urea on the activity over a range of substrate concentrations and investigate the detailed relation to conformational dynamics. We show that the increase in enzymatic activity in the presence of sub-denaturing concentrations of urea is caused by the alleviation of AMP inhibition. This effect can be traced to two mechanisms: a decrease in the affinity of the enzyme for AMP and a shift in dynamics that favors the open conformation of AK’s domains. Importantly, in the absence of inhibitory concentrations of AMP, the activity of AK is not enhanced by urea and can even be reduced. These results point to a nuanced role of the combination of domain closure dynamics and substrate binding, likely employed by a multitude of enzymes to regulate their activity.

## Results

### Urea relieves AMP inhibition

To capture urea’s effect on the enzymatic activity of AK, we assessed its impact at concentrations up to 0.8 M on both the forward and reverse reactions using coupled activity assays (Fig. 1). For the forward reaction (ATP + AMP → ADP + ADP), we monitored enzymatic velocity as a function of AMP concentration (Fig. 1a), revealing the familiar inhibition of the enzyme at high concentrations. The effect of urea varied greatly depending on the AMP concentration. At high AMP concentrations, urea increased the turnover; specifically, at 5 mM AMP, the velocity was 1.6 times higher in the presence of 0.8 M urea, in agreement with the results of Zhang and coworkers.^29^ However, the opposite effect was observed when the AMP concentration was low (Fig. 1a), where the turnover diminished by 40% at 100 µM AMP, suggesting that urea activated only AMP- inhibited AK.

**Figure 1.**
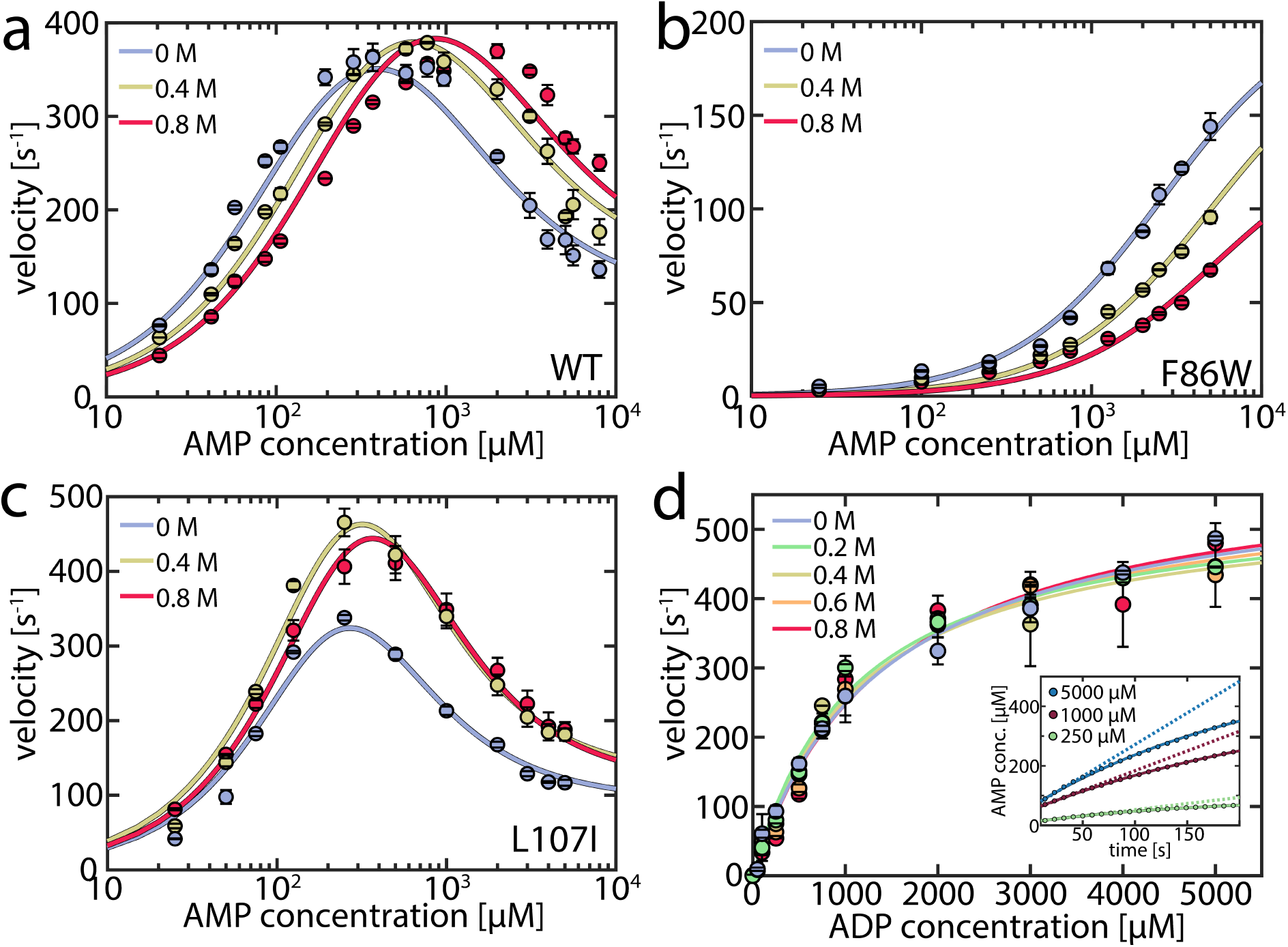
Enzymatic activity in the presence of urea. a) Enzymatic velocity of WT AK as a function of AMP concentration. The presence of urea (blue to red curves) gradually alleviates substrate inhibition at high AMP concentrations but inhibits AK at concentrations below 300-400 μM AMP. The solid lines are fits to the model depicted in Fig. 5a. Error bars indicate the standard error of the mean of at least 2 measurements. b) As in (a) for the non-inhibited mutant F86W. c) As in (a) for the strongly inhibited mutant L107I. d). Enzymatic velocity *v_0_* for the backward reaction at different urea concentrations. *v_0_* can be described by simple Michaelis-Menten kinetics and is hardly affected by the urea concentration. The inset shows the product inhibition of the backward reaction. Shown is the AMP concentration following the start of the reaction for representative initial ADP concentrations of 250 µM (green), 1000 µM (red) and 5000 µM (blue). AMP is formed tantamount to ATP and lowers turnover accordingly as the reaction progresses, resulting in a nonlinear formation of AMP. The solid lines are fits according to De La Cruz.^33^ The dashed lines represent the extrapolated product formation without product inhibition.

As a result, we anticipated that different effects would arise in mutant proteins based on their propensity for substrate inhibition. The F86W mutation, located in the AMP binding site, has been shown to eliminate substrate inhibition.^17, 32^ Urea did not enhance the enzymatic velocity of F86W at any substrate concentration (Fig. 1b). In contrast, the strongly inhibited mutant L107I^17^ exhibited a higher velocity in the presence of urea (Fig. 1c), similar to the wild type enzyme (WT).

A connection between AMP inhibition and activity enhancement by urea was also observed in the reverse reaction (ADP + ADP → AMP + ATP). In this direction, AMP acted as a product inhibitor, leading to a non- linearity in the steady-state time course (Fig. 1d inset). To extract the actual initial velocities *v_0_* for ADP conversion, we applied the approach of De La Cruz *et al.*^33^ and assessed activity using

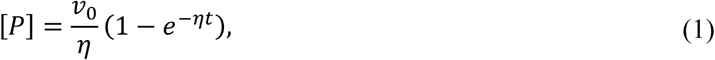

where [*P*] is the concentration of the steady-state enzyme-catalyzed product, while the factor *η* describes the non-linearity. The extracted *v_0_* values were fitted to the Michalis-Menten equation (Fig. 1d). Upon the addition of urea, neither the maximum velocity *v_max_* nor the Michaelis constant *K*_M_ changed significantly (Table S1). However, urea notably mitigated the inhibition by AMP, expressed in a reduced nonlinear behavior (Fig. S1a, Table S2). The parameter *η* decreased by up to 38±6% in the presence of urea, demonstrating urea’s propensity to reduce inhibition. Additional experiments in the presence of 10 mM AMP (Fig. S1b) further confirmed the role of AMP as a competitive inhibitor and that this inhibition is attenuated by urea. In conclusion, urea reduced AK’s inhibition by AMP but did not increase activity when this substrate/inhibitor was absent.

We hypothesized that urea could reduce AMP inhibition of AK in two ways. First, it could change the affinity of the enzyme to its substrate/inhibitor AMP, and second, it could affect the conformational dynamics of the enzyme. To probe the first factor, we measured AMP affinity by microscale thermophoresis (MST) across a range of urea concentrations. As shown in Fig. 2, urea reduced the affinity for AMP, resulting in a 2.3–fold increase in *K*_d_ (AMP) in the presence of 0.8 M urea. In contrast, the affinity for ATP was barely affected by urea.^30^

**Figure 2.**
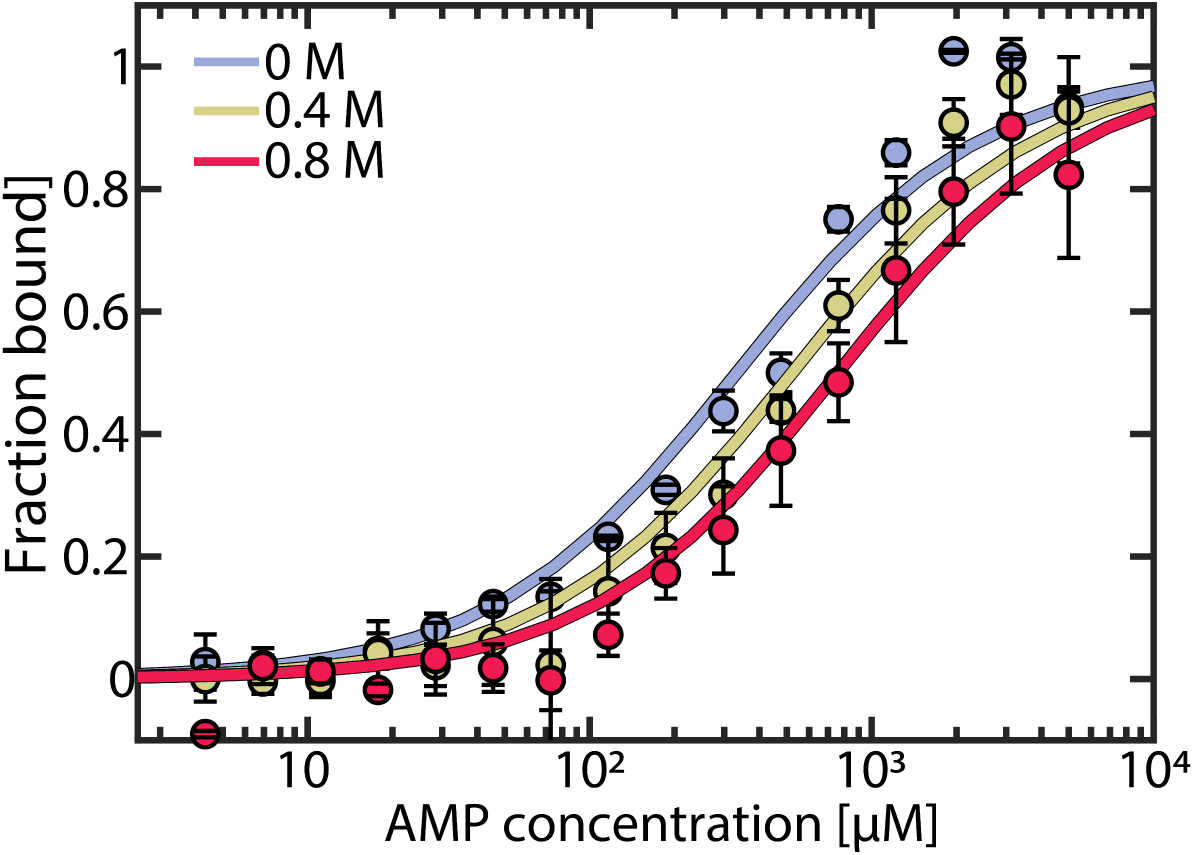
AMP binding to AK measured by MST. The fraction of bound AK as a function of AMP concentration was measured at urea concentrations of 0 M (blue), 0.4 M (yellow) and 0.8 M (red). Data was fitted to obtain *K*_D_ values of 332 ± 40 µM, 522 ± 58 µM and 758 ± 76 µM, respectively. Error bars indicate the standard errors of the mean of 3 measurements.

### Urea affects the conformational transitions of AK

To test whether urea further alters AK’s activity by affecting the conformational dynamics, we used smFRET spectroscopy to monitor the impact of urea on the enzyme’s domain motions at sub-denaturing concentrations. The double-labeled mutant A73C-V142C of AK was employed as an excellent probe for the LID-CORE distance (Fig. 3a).^16, 17^ We conducted smFRET experiments on freely diffusing enzyme molecules and calculated FRET efficiency histograms. In the absence of substrates, the histogram was found to peak at a value of ∼0.4, with a tail towards high values (Fig. 3b), demonstrating a prevalence of open conformations of the apoprotein. Upon the addition of urea, only minor changes in the FRET efficiency distribution were observed, at least at urea concentrations below 1 M (Fig. S2a+b). Within this concentration range, only a tiny fraction of protein molecules unfolded, as confirmed by CD spectroscopy (Fig. S3).

**Figure 3.**
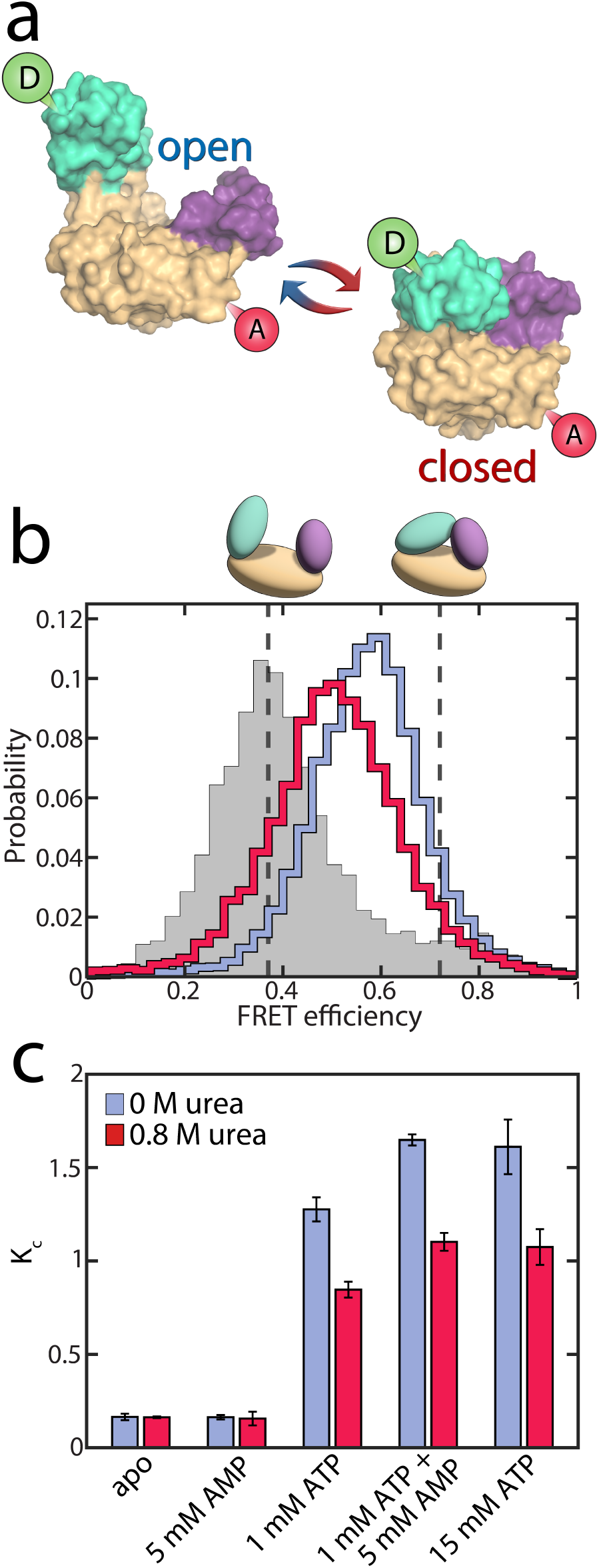
Conformational equilibrium in AK. a) Structure of AK with its three domains. The CORE domain (yellow) connects the LID domain (teal) and the NMP domain (purple). The spheres indicate the positions for the attachment of donor (green) and acceptor (red) dyes. The protein can undergo a conformational change from the open state (left, PDB: 4AKE) towards the closed conformation (right, PDB: 1AKE). b) The FRET efficiency histogram of the apoprotein is shown in grey. The solid lines correspond to histograms in the presence of substrates (1 mM ATP, 5 mM AMP and 417 µM ADP) with 0 M urea (blue) and 0.8 M urea (red). The dashed grey lines indicate the FRET efficiency values of the open (0.37±0.01) and closed (0.72±0.01) state obtained from the H^2^MM analysis. c) *K*_C_, which is the ratio of the two conformations in (a), as a function of substrate concentration. The occupancies of closed and open state were determined by H^2^MM and are shown in the absence (blue) and the presence of 0.8 M urea (red). Without ATP, urea does not affect *K*_C_ significantly. With ATP present and the LID domain predominantly closed, urea reduces the population of the closed state.

In the presence of nucleotide substrates, urea had a much more pronounced effect on the conformational distribution (Fig. S2c+d). ATP/ADP binding shifted the histogram towards higher FRET efficiencies (blue curve in Fig. 3b), corresponding to a reduced distance between the two dyes due to domain closure. A detailed analysis of the impact of substrate binding itself can be found in our recent publications,^16, 17^ and a summary is provided in the Supporting Information (Fig. S4). The addition of urea partially opposed the effects of substrate binding, shifting the histogram back towards lower FRET efficiencies (red curve in Fig. 3b).

Quantitative information on the protein dynamics underlying the FRET efficiency histograms was extracted by statistical analysis using the algorithm H^2^MM developed in our lab.^34^ Based on hidden Markov models, this algorithm can determine the populations of the open and closed states as well as their interconversion rates from a photon-by-photon analysis of single-molecule trajectories. In our analysis, the FRET efficiencies of the two states were optimized globally to provide the best fit across varying substrate and urea concentrations (*E*_open_=0.37±0.02, *E*_closed_=0.72±0.02). This procedure was employed since the structures of the two conformational states were considered unaffected by urea within the given concentration range, as indicated by the minimal changes in the FRET efficiency distribution of the apoprotein (Fig. S2a) and the fraction of unfolded protein measured by CD (Fig. S3). In contrast to their structure, the distribution of these conformational states was affected by urea. The H^2^MM analysis was validated using three procedures: a recoloring analysis (Fig. S5), a dwell-time analysis (Fig. S6, Table S3) and a time-resolved burst variance analysis (Fig. S7).^35^ Representative trajectories, including state assignment with the Viterbi algorithm,^34^ are shown in Fig. S8.

To characterize the changes in the population of the open (*P_o_*_pen_) and closed state (*P*_closed_), we used an effective equilibrium coefficient *K*_C_

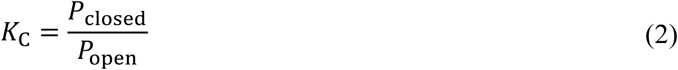

As already inferred from the analysis of the FRET efficiency histograms, the population of the closed, high FRET state of the enzyme was small under apo conditions (*K*_C_ = 0.16±0.02, blue bar in Fig. 3c) and remained similar in the presence of only AMP (*K*_C_ = 0.16±0.01). However, the closed state population increased strongly when ATP was present, with *K*_C_ = 1.28±0.06 in the presence of 1 mM ATP and *K*_C_ = 1.65±0.03 with 1 mM ATP + 5 mM AMP.^17^ The addition of 0.8 M urea changed the conformational-state populations, as shown by the red bars in Fig. 3c. In the absence of ATP, where the open conformation dominated, urea did not affect the population of the two states, as already discussed. In contrast, in the presence of ATP, urea significantly lowered *K*_C_ (*K*_C_ = 0.85±0.04 with 1 mM ATP + urea, and *K*_C_ = 1.10±0.05 with 1 mM ATP + 5 mM AMP + urea). This observation coincided with the findings of Rogne *et al.*, who proposed that the compact closed state is more susceptible to urea than the apoprotein, based on indirect observations of the conformational equilibria using NMR spectroscopy.^30^

The reduction in K_C_ persisted even at a very high ATP concentration of 15 mM. Given that this ATP concentration is 300 times higher than the dissociation constant *K*_D_ for ATP,^36^ we concluded that it is unlikely that the conformational shift is caused by a reduction of the ATP-bound species. This conclusion was in agreement with NMR spectroscopy, which suggested that 1 M urea has only a minor effect on *K*_D_ (ATP).^30^ Instead, urea influenced the interconversion rates between the open and closed states in the substrate-bound protein, favoring the open conformation of the protein (see below). We also examined the effect of urea on the two mutant proteins F86W and L107I: a similar reduction in *K*_C_ was observed for both mutants (Fig. S9), although they varied greatly in terms of their catalytic properties (Fig. 1b+c).

### LID domain opening is accelerated by urea

A great advantage of single-molecule studies is that they allow for direct access not only to the occupancy of each state but also to the specific rates for domain opening and closing. To establish a correlation between these rates and protein turnover, we examined the apparent closing and opening rates (Fig. 4) across increasing concentrations of AMP while maintaining a fixed ATP concentration of 1 mM. This setting resembled the conditions for the enzymatic activity assay, allowing us to assess how the pronounced substrate inhibition by AMP^32, 37^ affects the conformational dynamics. Domain closing (Fig. 4 upper panel) and opening rates (lower panel) were both on the microsecond time scale, two orders of magnitude faster than the enzyme’s turnover rate, which amounted to a maximum of ∼370 s^-1^ under these conditions. Nevertheless, activity and protein dynamics were closely linked: in the absence of urea (blue curves), we noted an increase in the LID closing rate when the AMP concentrations exceeded ∼ 500 μM, coinciding with a drop in enzymatic velocity due to the AMP inhibition (Fig. 1a). In contrast, domain opening was unaffected by AMP. In the presence of 0.8 M urea, at small AMP concentrations, both domain opening and closing were slowed by urea. Adding AMP accelerated domain closing, resulting in a similar closing rate at 0 and 0.8 M urea at high AMP levels. Interestingly, with urea present, the opening rate was *also* influenced by AMP, leading to an AMP-dependent acceleration similar to the closing rate. At 10 mM AMP, domain opening was 1.3±0.1 times faster in the presence of urea than in its absence.

**Figure 4.**
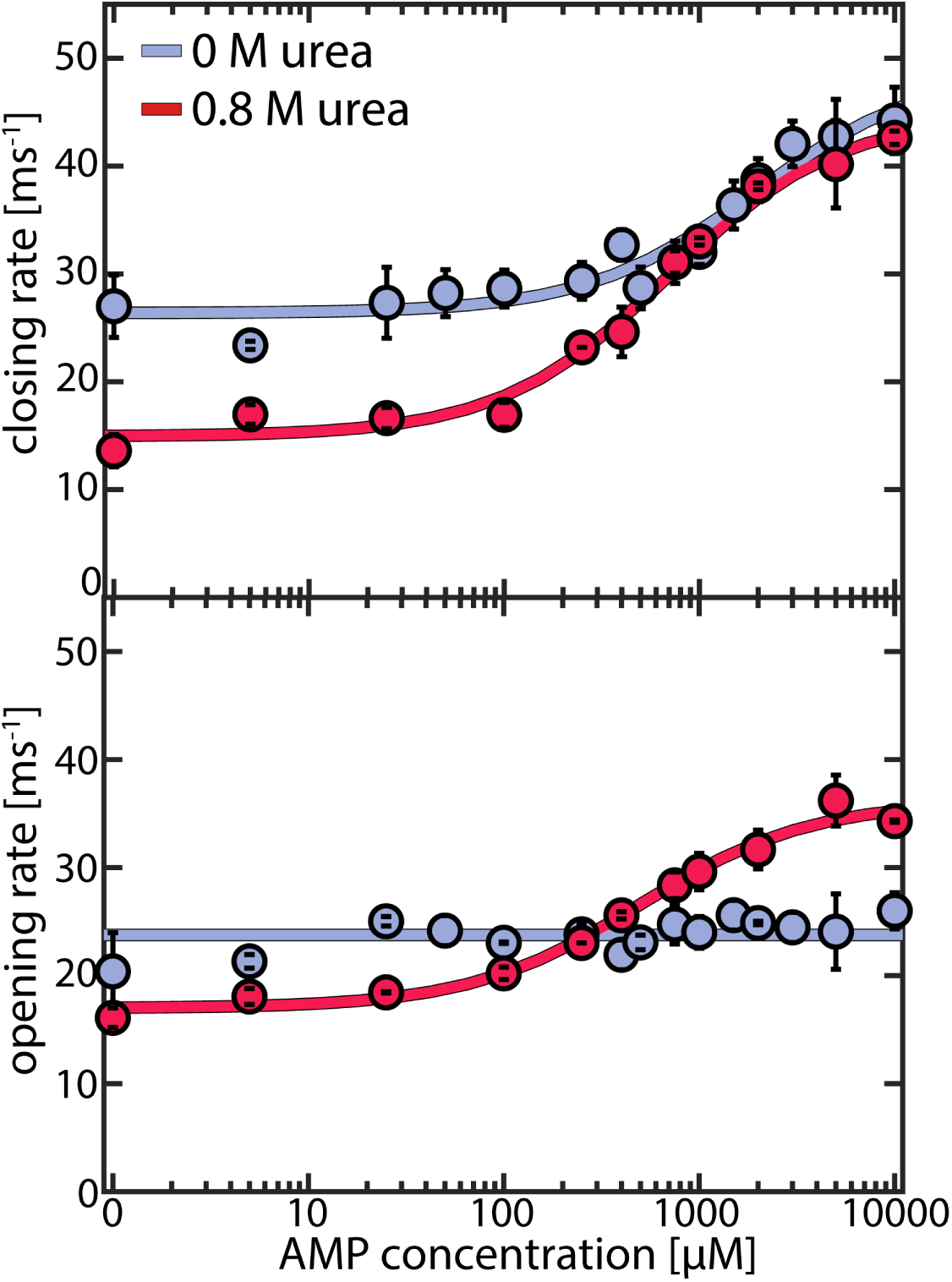
Effect of urea on AMP-dependent domain closing and opening rates. Closing (upper panel) and opening (lower panel) rates for the WT protein as a function of AMP concentration. Experiments were conducted at a fixed ATP concentration of 1 mM and appropriate concentrations of ADP to maintain equilibrium, as described in the Supplementary Methods (Table S4). Blue points are measurements without urea and red points in the presence of 0.8 M urea. Solid lines indicate fits to the model described in Supporting Note 2. Error bars indicate the standard error of the mean of at least 2 measurements.

To confirm our observations, we measured the effect of urea on the two mutant enzymes L107I and F86W. For L107I, which shows strong substrate inhibition,^17^ we observed very similar effects to the WT (Fig. S10). In contrast, for the non-inhibited mutant F86W^17, 32^ (Fig. S11), urea had a more pronounced impact on domain closing than on opening, as observed for the WT/ L107I only in the absence of inhibiting concentrations of AMP. This suggested that urea affects the inhibited and non-inhibited species differently.

## Discussion

### Fitting enzymatic velocity in the presence of conformational dynamics

The experimental observations described in previous sections revealed that urea significantly modulated conformational dynamics by favoring the open substrate-bound conformation. Urea also lowered the affinity for AMP. How could these effects lead to the surge in velocity observed at high AMP concentrations? To address this, we employed the kinetic model introduced in our previous work^17^ illustrated in Fig. 5a and fully explained in Supporting Note 1. In brief, this model accounts for conformational dynamics by attributing both open and closed conformations to each substrate-bound species, with specific transition rates between these states taken directly from the single-molecule experiments (refer to Supporting Note 2 and Table S5 for details). The catalytically active ternary complex ETM (enzyme + ATP + AMP) can be formed via two kinetically distinct pathways differing in the order of ligand binding.^38^ Starting from the apoenzyme E, either ATP or AMP can bind first, resulting in the species ET and EM, respectively. Subsequent binding of the corresponding co-substrate leads to two different ternary complexes: ETM_i_ (in the “ATP first” pathway) and EMT_i_ (in the “AMP first” pathway). These two states are catalytically inactive due to unsuitable substrate alignment. To achieve the catalytically active ETM configuration, the substrates have to undergo rearrangement, which is only possible in the open conformation.^25, 39^ If the rate constant for this process, *k*_r_, is slow in one of the pathways, this creates a kinetic barrier leading to substrate inhibition.

**Figure 5.**
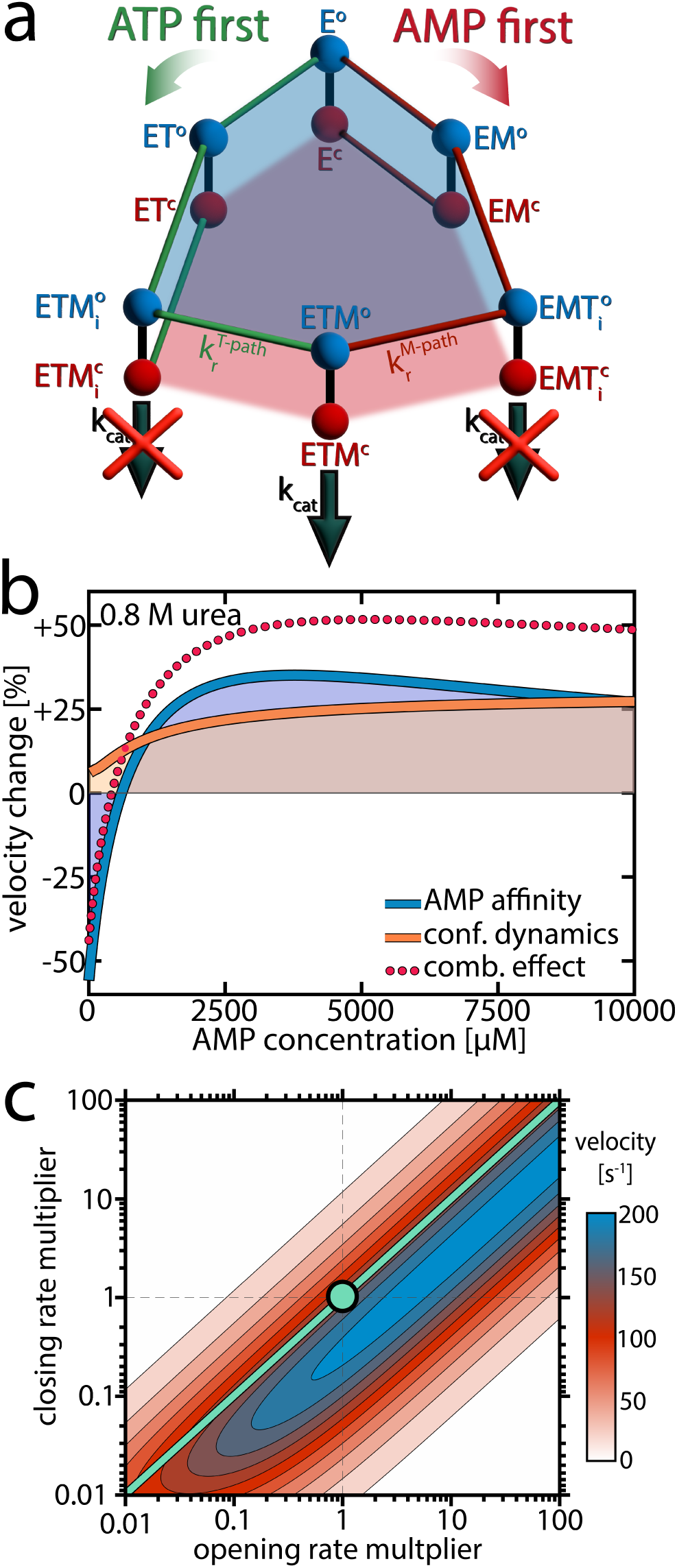
Explaining the effect or urea on enzymatic activity. a) A model for the enzymatic activity that considers six different substrate-bound species. Each state exists either in the open (o, blue) or closed (c, red) conformation.^25^ In the "ATP first" pathway (green), ATP binds to the apoenzyme (E) first, resulting in the ATP-bound state (ET). Alternatively, in the "AMP first" pathway (red), AMP binds first, leading to the AMP-bound state (EM). Subsequent binding of the respective co-substrate in both pathways produces a ternary complex. In the "ATP first" pathway, this leads to the inactive ETM_i_ state, while in the "AMP first" pathway to the inactive EMT_i_ state. To reach the catalytically competent ETM state, the substrates must rearrange into the native binding pose.^25^ The rate constant for this process, *k*_r_, depends on the binding order of the substrates. A slow 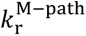 creates a kinetic barrier that invokes AMP’s substrate inhibition. An extensive description of the model is provided in Supporting Note 1. b) Based on the model in (a), we simulated the effect of urea on enzymatic velocity as a function of the AMP concentration. To obtain the blue line, we altered AMP affinity according to experimental observations, while preserving all other parameters, including those describing conformational dynamics, as determined at 0 M urea. In contrast, for the orange line, we altered the conformational dynamics while preserving AMP affinity. c) Enzymatic velocity as a function of the opening and closing rate. We simulated how the velocity changes when the opening and closing rates of the ATP-bound species (ET, ETM_i_, EMT_i_, ETM) are altered. The experimentally derived closing and opening rates were scaled by a factor between 0.01 and 100, at a substrate concentration of 10 mM AMP and 1 mM ATP. The turquoise circle denotes the experimental rates. Adjusting both opening and closing rates by the same factor does not change velocity, as indicated by the turquoise line, provided the rates are sufficiently large to avoid becoming rate-limiting.

We fitted this enzymatic model to the urea-dependent experimental data. Two sets of parameters were considered to be directly influenced by urea: the opening/closing rates measured in the smFRET experiments, and the AMP affinity values measured by MST. Other parameters in the fitting procedure, including the rate constants of phosphotransfer (*k*_cat_) and the substrate-rearrangement in the “ATP first” 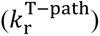 and “AMP first” 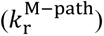 pathways, were considered unaffected by urea. This assumption was supported by the crystal structure of the closed state of AK, which indicated that the active site is well shielded from the solvent,^12^ rendering it less likely that urea directly affects the catalytic rate. The parameters were optimized globally across urea concentrations to match the enzymatic turnover as a function of either AMP or ATP concentration, with the results presented in Table 1 and Fig. 1. To estimate the confidence intervals for the fitted parameters, we monitored the increase in 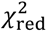 upon perturbing each parameter (Fig. S12-S14). The continuous lines in Fig. 1a-c show the fitted curves for the AMP-dependent turnover, and Fig. S15 shows the fitted curves for the ATP-dependent turnover. Overall, the model adequately captured all the nuanced features imposed by the addition of urea.

**Table 1.** Fit parameters describing the activity of AK.

| parameter |  | urea [M] | WT | F86W | L107I |
| --- | --- | --- | --- | --- | --- |
| rate constant<br>[s<sup>-1</sup>] | $k_{cat}$ | 0-0.8 | $3.1 \cdot 10^3$ | 492 | $\geq 5.3 \cdot 10^3$ <sup>d</sup> |
| | $k_r^{T-path}$ | 0-0.8 | $4.2 \cdot 10^3$ | $3.3 \cdot 10^3$ | $7.9 \cdot 10^3$ |
| | $k_r^{M-path}$ | 0-0.8 | 370 | - <sup>c</sup> | 276 |
| affinity<br>[μM] | $K_d$ (ATP) | 0-0.8 | 50 <sup>a</sup> | 62 | 10 |
| | $K_d$ (AMP) | 0 | 332 <sup>b</sup> | $1.3 \cdot 10^3$ | 628 |
| | | 0.4 | 522 <sup>b</sup> | $2.8 \cdot 10^3$ | 681 |
| | | 0.8 | 758 <sup>b</sup> | $4.9 \cdot 10^3$ | 890 |

<sup>a</sup> taken from Rogne *et al.*<sup>30</sup>
<sup>b</sup> determined by MST
<sup>c</sup> not obtainable due to lack of substrate inhibition
<sup>d</sup> only a lower bound could be determined reliably

### A lower affinity for AMP favors the more productive “ATP first” pathway

Notably, the observed changes in the parameters describing conformational dynamics and AMP affinity were sufficient to account for the pronounced velocity changes. But how can we explain these changes, and to what extent do the individual effects contribute to the overall change in activity? We tested computationally how the enzymatic velocity would change when only one of the parameters is affected by urea at a concentration of 0.8 M. As shown in Fig. 5, this analysis allowed us to isolate the contributions of each parameter.

First, we considered the scenario where AMP affinity was reduced while all other parameters, including those describing conformational dynamics, remained as determined at 0 M urea. The outcome is depicted by the blue line in Fig. 5b: At low AMP concentrations, the velocity decreased compared to the situation without urea, but at intermediate concentrations the velocity actually increased. To understand these opposing phenomena, it is important to observe the role of AK’s two competing mechanistic paths (Figure 5a). At low AMP concentrations, where initial AMP binding is unlikely, only the effective “ATP first” path is significantly populated. Therefore, reducing AMP affinity by the addition of urea essentially diminishes the flux through this path, with a strong detrimental effect on turnover.

In contrast, when the AMP concentration gets larger, both pathways are populated and the distribution between them determines the overall turnover. Lowering AMP affinity reduces the likelihood for initial AMP binding, shifting the population away from the unproductive “AMP first” path. As the rearrangement rate constant 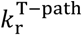 is much higher than 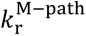 (Table 1), the velocity gain in the “ATP first” path outweighs the loss in the “AMP first” path. Finally, at very high AMP concentrations that far exceed *K*_D_ (AMP), reducing AMP affinity has no significant effect anymore. A more detailed analysis of how the AMP affinity affects the flux in the two pathways is given in Fig. S16.

### A conformational equilibrium determines AK’s activity

We next isolated how the modulation of conformational dynamics by urea affects enzymatic velocity. The orange curve in Fig. 5b represents how the velocity changed in response to a reduction in the closed-to-open ratio *K*_C_ for different substrate-bound species (while preserving nucleotide affinity). Promoting the open conformation enhanced activity, especially at high AMP concentrations where the “AMP first” pathway was predominantly occupied. This can be understood by recognizing that in the “AMP first” path *k*_r_ is much slower than *k*_cat_ (Table 1), making substrate rearrangement the rate-limiting step. Although shifting the closed-to-open ratio towards the open state slightly impeded the catalytic step, it substantially facilitated the rate-limiting substrate rearrangement, which was much more impactful.

In Fig. 5c we simulated how turnover changed when the balance between the open and closed states was altered. For this simulation, we multiplied either the opening or closing rates of the ATP-bound species by a factor ranging from 0.01 to 100, while keeping all other parameters as determined experimentally at 0 M urea. The substrate concentrations used in this calculation were 10 mM AMP and 1 mM ATP, meaning the “AMP first” path was primarily occupied, and substrate rearrangement was slow. Thus, reducing *K*_C_ by increasing the opening rate and/or reducing the closing rate enhanced enzymatic velocity. Adjusting both opening and closing rates by the same factor did not change velocity, as indicated by the turquoise line, provided these rates remained fast enough not to impede the rate-limiting step.

This calculation shows that the optimal balance between open and closed states depends on the rate constants

*k*_r_ and *k*_cat_. For AK, where 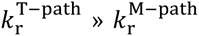, this means that the optimum also depends on substrate concentrations, as they determine the population of the two competing pathways. Interestingly, when the simulation was conducted under physiological concentrations of the nucleotides in *E. coli* (∼5 mM ATP, 300 μM AMP),^40^ the optimal open/closed ratio obtained in the calculation was close to the experimental *K*_C_ value (Fig. S17), suggesting the enzyme has evolved to optimize *K*_C_ for maximal turnover under physiological substrate concentrations.

In summary, at low AMP concentrations the reduction of AMP affinity by urea leads to a loss of activity. At intermediate AMP concentrations, reducing AMP affinity has a positive effect as it increases the population of the much more active “ATP first” path. Further, at high AMP concentrations changes in the conformational dynamics become highly important, and facilitation of substrate rearrangement by the modulation of the conformational equilibrium becomes decisive.

## Conclusion

### Reducing substrate affinity can rescue an enzyme from an inefficient pathway

The activating effect of urea defies initial expectations. Not only can a denaturant enhance enzymatic activity, but it does so by reducing substrate affinity and by promoting a non-catalytically active conformation. However, the model shown in Fig. 5a comprehensively explains these effects. The first important aspect is that the urea-induced reduction of AMP affinity allows for a faster conversion into the catalytically active species by promoting a kinetically faster pathway. This concept aligns with theoretical insights, which suggested that enzymatic reactions can be accelerated by increasing the rate of substrate unbinding when the enzyme occasionally gets trapped in an unproductive intermediate.^41, 42^ In such cases, accelerating substrate escape from the unproductive intermediate paradoxically enhances enzymatic velocity.^42^ In our model, the “AMP first” path represents such a trap, as the rate towards the catalytically active ETM species is much lower than in the “ATP first” path. Decreasing the affinity for the substrate AMP can expedite catalysis by lowering the chances of entering a less productive pathway, even though the overall fraction of time the enzyme is substrate-bound is reduced. However, when the amount of enzyme trapped in the low-efficiency pathway is marginal, as at low AMP concentrations, reducing AMP affinity does not accelerate catalysis but rather impedes it.

Activation by denaturants has also been reported for other enzymes.^43–46^ For instance, in human biliverdin- IXα reductase urea induces the formation of a partially unfolded intermediate. This intermediate follows a kinetically distinct path that bypasses the potent substrate inhibition by biliverdin.^43^ Kinetically distinct pathways may exist in many different enzymes,^47^ and controlling the flux through these pathways is a potent tool for manipulating protein activity.

### Favoring the open conformation facilitates productive substrate binding

The second crucial factor for successful catalysis is the correct positioning of the substrates. Substrate mobility is significantly restricted when the enzyme is closed, preventing reorientation in the closed conformation.^25, 39^ Urea promotes the open conformation of AK (Fig. 3c) and thus assists the reorientation process. Experiments utilizing a fluorescent substrate analog binding at the ATP binding site of AK suggested that urea indeed increases substrate flexibility at the active site.^29^ Enhanced flexibility and fast positioning of the substrates might be essential for many enzymes to express their full catalytic potential.^48^ Evidence for this hypothesis is found in other examples of denaturant-induced activation, such as in prostaglandin D synthase^46, 49^ and dihydrofolate reductase.^44, 45^ In both cases, the increase in activity has been attributed to enhanced flexibility at the active site.

However, promoting open conformations is not always beneficial for enzymatic activity. The optimal value for *K*_C_ depends on the rate constants for substrate reorientation and phosphotransfer. If *k*_r_»*k*_cat_, as in the non-inhibited mutant protein F86W (Table 1), phosphotransfer constitutes the rate-limiting step. Lowering *K*_C_ limits the fraction of molecules available for phosphotransfer and is thus detrimental to turnover (Fig. 1b).

In *E. coli*, under physiological concentrations of nucleotides^40^, the “ATP first” path predominantly drives the flux through the enzyme. With 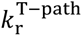 approximately equal to *k*_cat_, optimal turnover is achieved when open and closed conformations are sampled to a similar extent. Experimental *K*_C_ values close to ∼1 suggest that AK has evolved to sustain this optimal equilibrium, ensuring that both substrate reorientation and phosphotransfer occur at comparable rates under physiological conditions.

### Fast Dynamics Governing Slow Biological Functions

An important prerequisite for controlling activity via *K*_C_ is that the conformational dynamics must be faster than the associated processes of substrate reorientation and catalysis. Otherwise, domain opening and closing become rate-limiting. As demonstrated in Fig. 5c, when the opening and closing rates approach the timescale of catalysis/rearrangement, a significant decrease in turnover is observed even if *K*_C_ is preserved. Essentially, the protein activity, which occurs on a millisecond time scale, is governed by conformational dynamics on a much faster, microsecond time scale. These motions on two different time scales influence each other and are crucial for the protein’s overall function. While the fast conformational equilibrium determines the occupancy of active and inactive states, substrate binding alters the free energy landscape and the energy barriers between these states. Interestingly, this behavior has recently been observed in several other proteins,^50, 51^ such as enzymes,^52, 53^ membrane proteins^54^ and large disaggregation machines.^55^ Thus, a growing body of evidence points towards a general, *two-time-scale paradigm* for biological machines: fast equilibrium fluctuations occur on the microsecond-millisecond time scale, while on a slower time scale, free energy is invested to enable functional transitions.^56^

### The physiological role of the open/closed ratio

Our previous study demonstrated that substrate inhibition is an evolutionary well-conserved feature in AK.^17^ The strength of substrate inhibition appears to be optimized to ensure maximal bacterial fitness.^57^ Consequently, manipulating the conformational equilibrium in AK by any means directly alters its activity, potentially affecting the entire organism. Despite urea’s perturbing effect on protein structure and dynamics, some species accumulate urea as the primary osmolyte.^58^ For instance, urea concentrations in the kidney of a xeric desert rodent can reach 3-4 M during extreme water stress.^59^ In cartilaginous fish and the coelacanth, urea concentrations can be as high as 0.4 M.^60^ To cope with such high urea levels, these fishes simultaneously accumulate a second set of nitrogenous osmolytes, namely methylamines such as trimethylamine N-oxide (TMAO).^58^ Contrary to urea, TMAO increases the stability of proteins; in the case of AK, it was shown to promote the more compact closed structure.^61, 62^ In line with the conclusions from our model, such a conformational change reduces the activity of the protein.^62^

The results in this paper underscore the importance of controlling the balance between different conformational states for the activity of enzymes. smFRET allows us to study not only the impact of urea on the conformational equilibrium but also the explicit rates for domain opening and closing. Insights from such studies can be harnessed to alter enzymatic properties. Targeted engineering of conformational dynamics to change enzyme selectivity and activity is currently still very challenging, but not impossible, as a number of literature examples show.^63^ Investigating the role of (very fast) conformational variation is indispensable for the rational design of enzymes with desired properties and to our understanding of enzyme catalysis.

## Material and Methods

### Protein expression, purification and labeling

Protein expression, purification and labeling were performed according to published protocols.^16, 17^ A summary is given in the Supporting Information.

### Enzymatic activity

Enzymatic assays in both directions were adapted from Pan *et al.*^64^ and performed on the labeled enzyme at urea concentrations between 0-0.8 M. Within this denaturant concentration range, the coupled enzymatic systems remain active (Fig. S18) and the fraction of unfolded AK itself did not increase, as confirmed by circular dichroism (CD) spectroscopy (Fig. S3). The velocity of the forward reaction (MgATP+AMP → MgADP+ADP) was monitored through the oxidation of NADH at 340 nm in coupling with pyruvate kinase and lactate dehydrogenase. The assay mixture was: 0-0.8 M urea, 4 nM AK, 50 mM Tris-HCl (pH 8.0), 100 mM KCl, 4 mM phosphoenolpyruvate, 5.0 mM MgCl_2_, 0.2 mM NADH, 10 units/mL pyruvate kinase, 15 units/mL lactate dehydrogenase, 0.25 mg/mL bovine serum albumin, 1 mM ATP and varying concentrations of AMP. The initial velocity was obtained by linearly fitting the NADH signal as a function of time.

The velocity of the backward reaction (MgADP+ADP → AMP +MgATP) was measured by following the reduction of NADP^+^ at 340 nm in a coupled reaction with hexokinase and glucose-6-phosphate dehydrogenase. The assay mixture was: 0-0.8 M urea, 4 nM AK, 50 mM Tris-HCl (pH 8.0), 100 mM KCl, 6.7 mM glucose, 0.67 mM NADP, 10 units/ml hexokinase, 10 units/ml glucose-6-phosphate dehydrogenase,

0.25 mg/ml bovine serum albumin, 10 mM MgCl_2_ and varying concentrations of ADP. The reaction rate decreased during the time course of the reaction due to the formation of the product and inhibitor AMP, resulting in a nonlinear increase in product concentration. The initial velocity *v_0_* at each concentration was obtained according to Eq. (2) and fitted to the Michaelis-Menten equation to obtain the maximum velocity *v*_max_ and the Michaelis constant *K*_M_.

### smFRET data acquisition and data analysis

The sample preparation for the single-molecule measurements is described in the Supporting Information. For data acquisition, a Microtime 200 system (PicoQuant) was used. Measurements were conducted in pulse-interleaved excitation mode,^65^ using a sequence of one pulse for acceptor excitation (564 nm) and three pulses for donor excitation (488 nm) at 40 MHz. The laser beams were focused 10 µm deep into the sample solution. Molecules diffusing through the focus created short bursts of photons that were divided into two channels according to their wavelengths, using a dichroic mirror (zt594rdc; Chroma) and filtered by band-pass filters (HC520/35 (Semrock) for the donor channel and ET 674/75m (Chroma) for the acceptor channel). Arrival times of these photons were registered by two single-photon avalanche photodiodes (SPCM-AQRH-14-TR, Excelitas) coupled to a time-correlated single-photon counting module (PicoHarp 400, PicoQuant).

To identify fluorescent bursts, the data was first smoothed with a running average of 15 photons. A cut-off time of 5 μs between individual photons was used to define each burst’s start and end points. Only fluorescence bursts with 50 photons or more were selected for further analysis. From each measurement, ∼10,000 burst events were collected. The raw FRET efficiency of each burst was calculated based on the photons detected in both channels following donor excitation only. The raw stoichiometry was obtained from the detected photons in both channels after both excitations, as described elsewhere.^66, 67^ A 2D histogram of raw stoichiometry versus raw FRET efficiency was generated, from which we extracted the amount of emitted donor photons leaking into the acceptor channel and the level of direct excitation of the acceptor dye by the 485 nm laser.^66^ The photon counts in both channels were corrected based on the calculated factors.^66, 68^ To obtain a corrected FRET histogram without the donor and acceptor-only populations, we selected only photon bursts with a stoichiometry corresponding to molecules with both donor and acceptor dyes.

The dynamics hidden in the FRET efficiency histograms were extracted using the H^2^MM algorithm.^34^ Only double-labeled molecules and photons arising from donor excitation were taken for this analysis. The FRET efficiency values of the open and closed states were optimized globally to give the best fit across varying substrate concentrations. This procedure was employed since the structures of the two states were considered unaltered by substrate binding. The transition rates between the states were optimized for each concentration individually. For measurements in the presence of urea, the FRET efficiency values of the open and closed states were fixed to the values obtained by the global analysis. At the same time, other parameters were optimized for each measurement individually.

### Fitting enzymatic activity

Parameters of the enzyme kinetic model shown in Figure 5a and described in Supporting Note 1 that could not be measured from studies of conformational dynamics and nucleotide affinities were obtained by a global fitting procedure. An extensive description of the procedure is given in the Supporting Information. In brief, we used experimentally observed opening and closing rates of the LID domain for the different substrate-bound species (Table S5) as inputs. For the WT, experimental substrate affinities were provided as inputs. The dissociation constant of AMP, *K*_d_ (AMP), was determined by MST as a function of urea concentration (Fig. 2), while the dissociation constant of ATP, *K*_d_ (ATP), was taken from Rogne *et al*. and considered as not significantly impacted by urea.^30^ Additional free parameters were the rate of the phosphotransfer step (*k*_cat_) and the rates of correct substrate positioning (*k*_r_) in the “ATP first” and “AMP first” pathways. All free parameters were optimized globally across urea concentrations by numerically solving a set of differential equations describing the kinetics as detailed in Supporting Note 1. In particular, we used MATLAB’s^69^ ordinary differential equation solver ode15s and performed a χ^2^-minimization of *v_0_* versus substrate concentration.

## ASSOCIATED CONTENT

### Supporting Information

The Supporting Information contains two Supporting Notes, detailing our model for the substrate inhibition and the analysis of opening and closing rates. The Supporting Methods contain a detailed description of the methodology. Additional information is provided in terms of Supporting Figures S1– S18 and Supporting Tables S1–S6 as well as supporting references.

### NOTES

The authors declare no competing financial interests.

## ACKNOWLEDGMENT

We thank Prof. Amnon Horovitz from the Weizmann Institute of Science for carefully reading and commenting on the manuscript. The work of G.H. was supported by a grant from the European Research Council (ERC) under the European Union’s Horizon 2020 Research and Innovation Program (grant agreement No 742637, SMALLOSTERY) and a grant from the Israel Science Foundation (no. 1250/19). The work of D.S. was supported by Deutsche Forschungsgemeinschaft (DFG, German Research Foundation, Projektnummer 490757872). The work of E.I.S. was supported by the NIH grant 5R35GM139571.

## AUTHOR CONTRIBUTIONS

D.S. conceived the work, acquired, analyzed and interpreted the data and wrote the paper; D.L. performed experiments; R.C. contributed to the interpretation of the data and writing of the paper; I.R. performed experiments, as well as contributed to the interpretation of the data and writing of the paper; H.M. contributed to the interpretation of the data; G.H. conceived and supervised the work and contributed to the interpretation of the data and writing of the paper.

