## Supplementary Information for "Enzyme activation by urea reveals the interplay between conformational dynamics and substrate binding: a single-molecule FRET study"

David Scheerer^a^, Dorit Levy^a^, Remi Casier^a^, Inbal Riven^a^, Hisham Mazal^a,b^, Gilad Haran^a*^

^a^ Department of Chemical and Biological Physics, Weizmann Institute of Science, Rehovot 761001, Israel

^b^ Max Planck Institute for the Science of Light, Erlangen 91058, Germany

**Supporting Note 1: Model for substrate inhibition by AMP**

Given that AMP inhibits AK's enzymatic activity and affects domain-closure dynamics, we postulated a correlation between these two phenomena (Scheme 1).^1^ Our model is based on the hypothesis that different closed states can be sampled depending upon the order of ligand binding. Some of these states are incompatible with catalysis and must transition to a catalysis-prone state before the reaction occurs. Regardless of ligand binding, the enzyme can always exist in both open (blue) and closed (red) states. States are defined only for the ATP-binding LID domain. The rates for the interconversion between the states are defined for domain opening as *k*_o_ and for domain closing as *k*_c_. Each substrate-bound species has specific opening and closing rates, designated by the superscript. The binding of AMP to the NMP domain is unaffected by the LID domain's status, allowing both open and closed states to bind AMP. However, ATP can bind only to the open state.

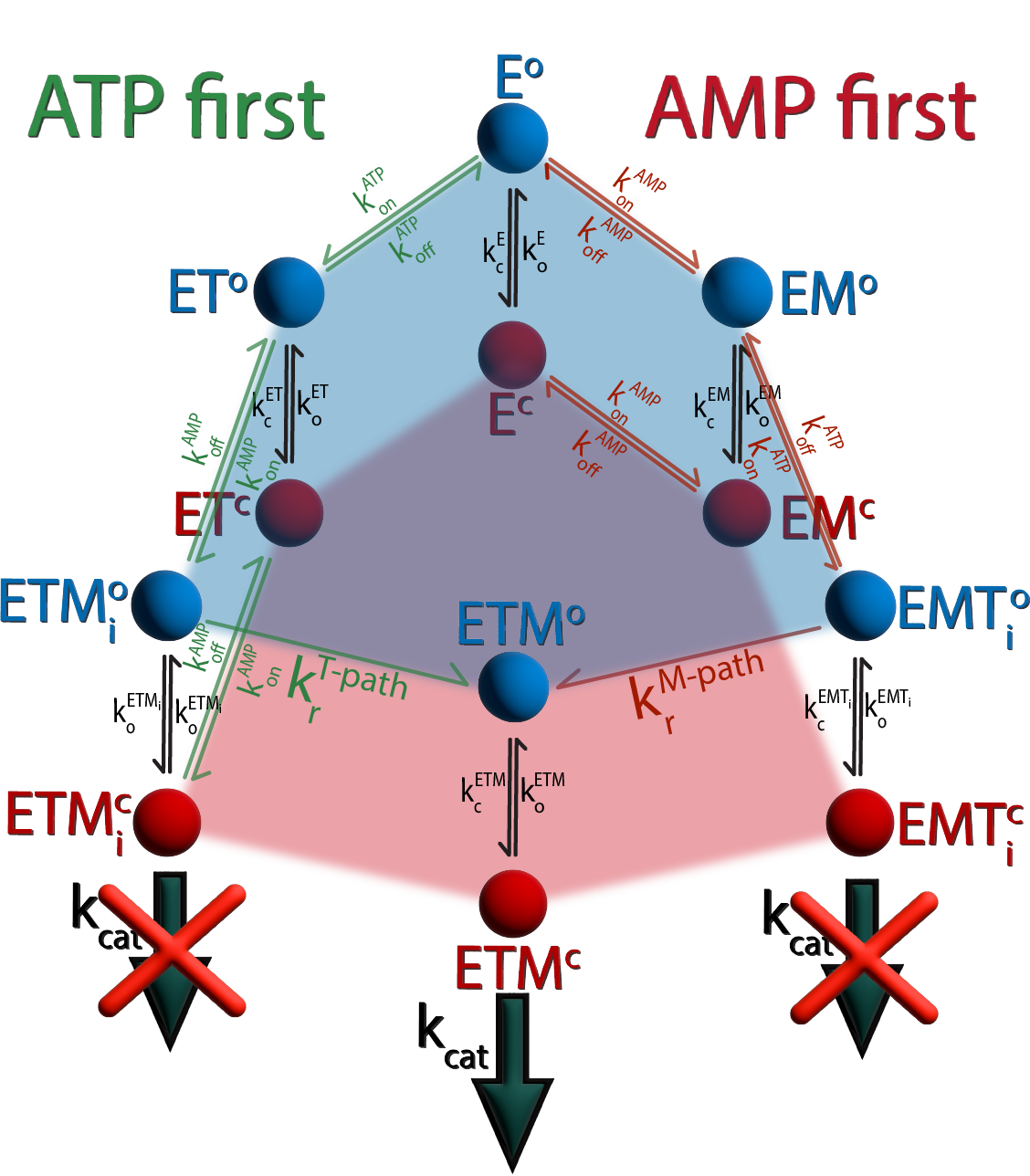

**Scheme 1.** Mechanism for the substrate inhibition in AK, based on two competing pathways.

The model in Scheme 1 leads to two scenarios regarding the order of nucleotide binding to the apoenzyme E. ATP can only bind if the LID domain is in the open state (E^o^). Should ATP bind first, this results in the ATP-bound state ET. Both the open (ET^o^) and closed conformation (ET^c^) of ET can subsequently bind AMP, forming the double substrate-bound state ETM_­i_. The initial encounter complex ETM_­i_ can exhibit diverse substrate binding poses, often differing significantly from the native structure.^2^ Catalysis is contingent upon the substrates repositioning themselves into the correct pose, denoted as ETM^o^, a process governed by the rearrangement rate constant $k_{r}^{T-path}$.

Conversely, if AMP binds first at the NMP domain, it yields the EM^o^ and EM^c^ complexes. The consequent binding of ATP leads to the formation of EMT_i_. Similar to ETM_­i_, we postulated that the substrates in this complex require rearrangement (with the rate constant $k_{r}^{M-path}$) to achieve the productive ETM state. ETM_i_ and EMT_i_ are distinguished by their binding order —ATP first, then AMP or vice versa— based on the hypothesis that initial AMP binding may result in a limited or different sampling of the nucleotide orientation.^3^ One reason might be that AMP binds in an orientation closer to the native binding pose when ATP is already bound and the LID domain is preferentially closed. A small rate constant $k_{r}^{M-path}$ effectively imposes a kinetic barrier within the "AMP first" path. The conversion of the inactive ETM_i_ and EMT_i_ states to the productive ETM state only occurs through their open states, as suggested recently by our MD simulations.^2^ The rates of phosphotransfer in the ETM state and subsequent product dissociation are combined in *k*_cat_. The rate constants for binding AMP or ATP are defined as $k_{\mathrm{on}}^{\mathrm{AMP}}$ and $k_{\mathrm{on}}^{\mathrm{ATP}}$, respectively. Instead of being converted to ADP, the substrates can also dissociate with the rates $k_{\mathrm{off}}^{\mathrm{AMP}}$ and $k_{\mathrm{off}}^{\mathrm{ATP}}$.

Essentially, the model introduces a kinetic barrier for the path in which AMP binds first. In a typical AK activity assay conducted under saturating ATP concentrations, the flux through the "ATP first" pathway predominates over that of the "AMP first" pathway at low levels of AMP (Fig. S17). However, as AMP levels rise, a larger fraction of the total flux diverts towards the "AMP first" path involving the unproductive ETM_i_ state, consequently diminishing the enzymatic velocity. Urea has the potential to influence both the conformational equilibrium and the affinity for AMP, thereby altering the flux in either of the two pathways.

**Supporting Note 2: Analysis of binding-state-dependent opening and closing rates**

The occupancy of the closed state in AK is dependent on the substrate concentration. Each substrate-bound species is presumed to have specific opening and closing rates. Our focus was primarily on the effect of AMP on the protein dynamics of the ATP-bound species. It was observed that minor AMP concentrations (<500 μM for the WT without urea) did not result in significant alterations in the opening and closing rates (Scheme 2a).

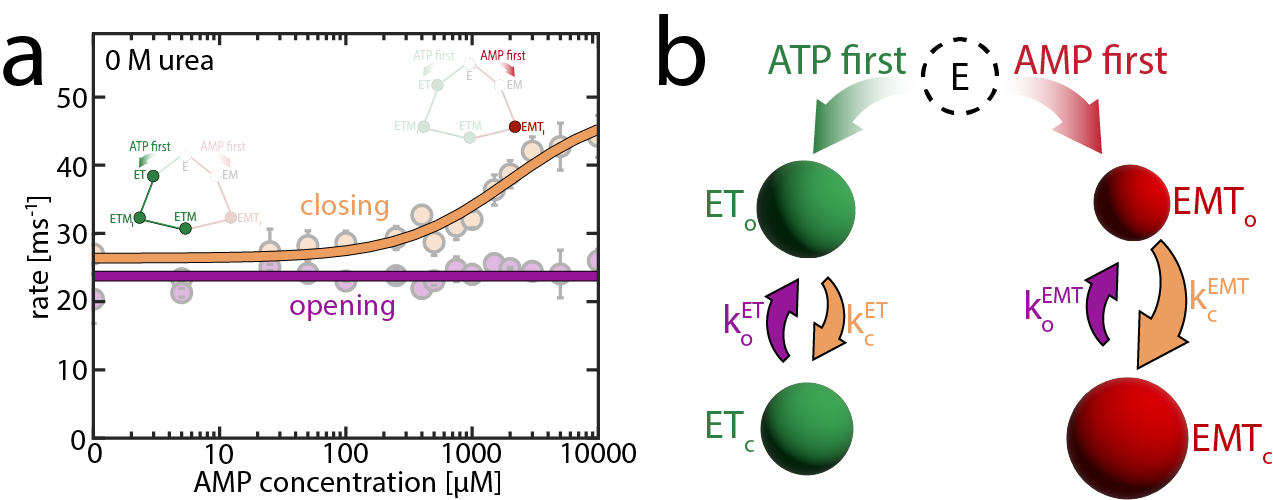

**Scheme 2.** Binding-state dependent opening and closing rates.

In this concentration range, the "ATP first" path is predominantly populated. Given the high enzymatic velocity at ~400 µM AMP (Fig. 1a main text), both the ETM_i_ and ETM state are likely populated to some extent. The absence of significant impact on the protein dynamics suggests that our single-molecule experiments cannot distinguish between the ET and the ETM/ETM_i_ species under these conditions. In Scheme 2b, these species are referred to as ET. In contrast, inhibitory concentrations of AMP above 500 μM increase the apparent closing rate (orange line in Scheme 2a), which we attribute to an increasing population of the "AMP first" pathway, particularly the inactive EMT_i_ state (denoted as EMT in Scheme 2b). In this scheme, we consider the enzyme as always bound to ATP, given that [ATP] » *K*_d_ (ATP). The different open enzyme species contribute to the apparent closing rate $k_{c}^{app}$ in proportion to their populations. The concentration of the more productive species ET/ETM/ETM^­^_i_ is denoted as $\left[ ET \right]$, while the concentration of molecules in the inactive EMT_­i_ species is denoted as $\left[ EMT \right]$, giving

$$k_{c}^{app}=\frac{\left[ ET \right]}{\left[ ET \right]+\left[ EMT \right]}\cdot k_{c}^{ET}+\frac{\left[ EMT \right]}{\left[ ET \right]+\left[ EMT \right]}\cdot k_{c}^{\mathrm{EMT}}=\frac{c_{50,\mathrm{AMP}}\cdot k_{c}^{\mathrm{ET}}+\left[ M \right]\cdot k_{c}^{\mathrm{EMT}}}{c_{50, \mathrm{AMP}}+\left[ M \right]},$$

(1)

with [*M*] as the AMP concentration, $k_{c}^{\mathrm{EMT}}$ as the closing rate of EMT_i_ species, $k_{c}^{ET}$ as the closing rate of ET/ETM/ETM^­^_i_. *c*_50,AMP_ is the AMP concentration at which both the ET and EMT species are equally populated.

$$c_{50,\mathrm{AMP}}=\frac{\left[ ET \right]\cdot\left[ M \right]}{\left[ EMT \right]}$$

(2)

Without urea, AMP does not significantly affect the apparent domain opening rate, so it was treated as constant over the complete AMP range (purple line in Scheme 2a). In contrast, an increase in the opening rate was detected with 0.8 M urea present (Fig. 4 main text). In this case, the opening and closing rates were globally optimized according to Eq. (1) and (3).

$$k_{c}^{app}=\frac{\left[ ET \right]}{\left[ ET \right]+\left[ EMT \right]}\cdot k_{o}^{ET}+\frac{\left[ EMT \right]}{\left[ ET \right]+\left[ EMT \right]}\cdot k_{o}^{\mathrm{EMT}}=\frac{c_{50,\mathrm{AMP}}\cdot k_{o}^{\mathrm{ET}}+\left[ M \right]\cdot k_{o}^{\mathrm{EMT}}}{c_{50, \mathrm{AMP}}+\left[ M \right]},$$

(3)

with $k_{o}^{\mathrm{EMT}}$ as the opening rate of EMT_i_ species and $k_{o}^{ET}$ as the opening rate of ET/ETM/ETM^­^_i_.

In the absence of ATP, the effect of AMP on protein dynamics is weak.^1^ Opening and closing rates for the EM species were measured in the presence of 5 mM AMP.

**Supporting Methods**

**Protein Expression and Labeling.** A pET15b expression vector containing the E. coli AK C77S gene with a six-residue histidine tag at the N terminus was used to express the protein for fluorescence experiments. Alanine-to-cysteine and valine-to-cysteine substitutions were introduced at positions 73 and 142 of the protein, respectively, by site-directed mutagenesis. The sequence of AK C77S/V142C/A73C was verified by DNA sequencing. The mutant was overexpressed by transformation into *E. coli* BL21 competent cells. The cells were lysed using high-energy sonication or a French cell press. The pellet was then separated from the protein-containing supernatant using high-speed centrifugation. The supernatant was separated on a Ni Sepharose column (GE Healthcare HisTrap HP). Fractions containing AK were pooled and run on a second gel filtration column (HiLoad 16/60 Superdex 75 prep grade; GE Healthcare) and eluted as a single peak. Labeling reactions were performed by first incubating protein samples with Alexa 594 maleimide (Invitrogen) at a molar ratio of 75%, separating labeled from unlabeled protein on a mono-Q 5/50 GL column (GE Healthcare), and later on labeling with an excess of Alexa 488 maleimide (Invitrogen).

**SmFRET measurements**

Single-molecule data was acquired on freely diffusing molecules using a Microtime 200 system (PicoQuant). Flow cells were prepared as described previously^4^ and filled with a mixture of 30 pM labeled enzyme, 50 mM Tris-HCl (pH 8.0), 100 mM KCl, 5 mM MgCl_2_, 0.01% Tween (Thermo Fisher), and substrates (ATP, ADP, AMP; Sigma). Importantly, we used ^31^P-NMR spectroscopy to verify that our ATP solutions contained no ADP. Substrate concentrations used for experiments under turnover conditions are given in Table S4. The appropriate ADP concentration to guarantee equilibrium (zero flux) was calculated using the following rate equation:

$$v\propto\frac{k_{1}\left[ M \right]\left[ T \right]}{K_{D,T}K_{D,M}}-\frac{k_{-1}\left[ D_{1} \right]\left[ D_{2} \right]}{K_{D,D1}K_{D,D2}},$$

(4)

where *k*_1_/ k_-1_ are the forward and backward rate constants for the reaction and [M] and [T] are the AMP and ATP concentrations. *K*_D,S_ is defined as the dissociation constant of the respective substrate (T,M,D_1_,D_2_) and values were taken from Sheng *et al..*^5^ [D_1_] and [D_2_] are the concentrations of ATP bound to the LID and NMP domain, respectively. [D_2_] was calculated using

$$\left[ D_{2} \right]=\frac{\frac{1}{K_{Mg}}+\left[ D \right]+\left[ Mg \right]-\sqrt{\left( \frac{1}{K_{Mg}}+\left[ D \right]+\left[ Mg \right] \right)^{2}-4\cdot\left[ D \right]\cdot[Mg]}}{2},$$

(5)

where [Mg] and *K*_Mg_^5, 6^ are the concentration and dissociation constant for magnesium, respectively. FRET efficiency histograms from the first and last 1 h of each measurement were shown to overlap, validating that the substrate concentration did not change during the measurement.

**Recoloring analysis**

We performed a recoloring analysis to verify the parameters obtained from the H^2^MM analysis.^4, 7^ In this method, the arrival times of photons in each data set are retained, but their "colors" (i.e. whether they belong to the donor or acceptor) are erased. A stochastic simulation based on the H^2^MM parameters is then used to reassign the photons to the two experimental channels, and FRET efficiency histograms are reconstructed. A good match between the original and recolored histograms indicated a successful analysis.

**Dwell-time analysis**

The dwell-time analysis yielded the distributions of times the protein spends in each state (in this case, the open and closed state). Here, we computed these distributions using a likelihood-weighted segmentation algorithm developed in-house.^8, 9^ In this analysis, for each burst, every possible sequence of states contributes a fraction of a count to each dwell time, equal to the likelihood of the sequence. In contrast with the more common dwell-time analysis based on the Viterbi algorithm, all possible state sequences were considered, not only the most likely one. A good agreement of rates obtained directly from the H^2^MM analysis and those obtained from the dwell-time analysis was taken as a validation of the analysis (Table S3).

**Time-resolved burst variance analysis (trBVA)**

To further validate the presence of dynamics on a µs-time scale, we employed trBVA, a recently published time-resolved version of burst variance analysis that can quantify kinetic rates at microsecond to millisecond timescales.^10^ Bursts were partitioned into segments with a fixed number of photons. The FRET variance was computed from these segments and compared with the variance expected from shot noise. Systematically varying the segment size can capture dynamics at different timescales. For this analysis, we utilized the same bursts as for H^2^MM. To minimize the impact of photophysical artifacts, we removed windows where consecutive photons of the same color are more than 30 µs apart. This allowed us to fit the data to a two-state model.^10^

**Preparation of urea solutions**

Solutions for measurements in the presence of urea were prepared immediately before the experiment. The urea concentration was determined by measuring its index of refraction using a Fisher-Abbe refractometer (Fisher Scientific Co.), according to the equation below:^11^

$$\left[ \mathrm{urea} \right]=117.66\cdot\Delta n+29.753\cdot{\Delta n}^{2}+185.56{\cdot\Delta n}^{3},$$

(6)

where $\left[ \mathrm{urea} \right]$ is the concentration of urea, and $\Delta n$ is the difference between the index of refraction of buffer with and without urea. Due to the high ionic strength, the pH of the urea solutions had to be corrected by using the equation below:^12^

(7)

$${pH}_{\mathrm{real}}={pH}_{\mathrm{app}}\left( -0.394\cdot\left[ urea \right]+0.06015{\cdot\left[ urea \right]}^{2}-7.157\cdot{10}^{-3}\cdot\left[ urea \right]^{3}+3.382\cdot{10}^{-4}{\cdot\left[ urea \right]}^{4} \right),$$

where ${pH}_{\mathrm{real}}$ is the corrected pH and ${pH}_{\mathrm{app}}$ the apparent pH measured by the pH-meter.

**Circular dichroism spectroscopy**

The unfolding status of the protein was monitored by the circular dichroism (CD) signal at 222 nm (Fig. S3). In the concentration range used for the enzymatic essays and smFRET experiments (0-0.8 M urea), the secondary structure content of AK does not change. The solid lines in the figure indicate fits to a sigmoidal function with flat baselines:^13, 14^

$$\theta_{222nm}=\frac{\theta_{N}+ \theta_{U}\cdot K_{\mathrm{obs}}}{1+K_{\mathrm{obs}}},$$

(8)

where *θ*_N_ and *θ*_U_ is the molar ellipticity of the native and unfolded state, respectively. *K*_obs_ is given by:

$$K_{\mathrm{obs}}=\exp\left( \frac{-{\Delta G}_{\mathrm{fold}}^{0}-m*\left[ urea \right]}{R*T} \right),$$

(9)

with ${\Delta G}_{\mathrm{fold}}^{0}$ the folding free energy difference without urea, the m-value being the proportionality constant for the effect of urea concentration, *R* the gas constant and *T* the absolute temperature.

**Microscale thermophoresis**

AK–AMP interactions were measured using the microscale thermophoresis (MST) instrument Monolith NT.115 (Nano-Temper Technologies). A series of 10 μL solutions at different substrate concentrations was prepared. Each solution was mixed with a 10 μL solution containing AK molecules labeled with Alexa488 to obtain a final AK concentration of 30 nM and loaded into a special coated capillary supplied by Nano-Temper. Fluorescent molecules were excited with a blue laser (470 nm, 50% LED power) to monitor the spatial distribution of molecules in the capillary. Thermophoresis was measured in each capillary by locally heating a defined sample volume with a focused infrared laser (40% MST power) for 30 s. Protein molecules diffused away from the peak of the temperature gradient formed by the laser. Bound and unbound molecules responded differently, which led to a different steady-state spatial distribution of fluorescence. The change in depletion in the presence of substrate was plotted and used to calculate the bound protein fraction. The dissociation constant was then obtained by fitting the results to a binding isotherm.

**Fitting enzymatic activity**

To fit enzymatic velocity based on the kinetic model described in Supporting Note 1: "Model for the substrate inhibition by AMP", we used experimentally observed opening and closing rates of the LID domain for the different substrate-bound species as inputs. These rates were determined as described in Supporting Note 2: "Analysis of binding-state dependent opening and closing rates" and are listed in Table S5. The protein dynamics of the ET, ETM_i_ and ETM species were assumed to be the same, as minor concentrations of AMP (< 1000 μM) did not affect the opening and closing rates (Fig. 4, main text). The open and closed states are defined only for the ATP-binding LID domain. It was assumed that ATP can bind only to the open state, while AMP can bind to both open and closed states, as it binds to the NMP domain.

For the WT, experimental substrate affinities were provided as an input. The dissociation constant of AMP, *K*_d_ (AMP), as a function of urea concentration was determined by MST (Fig. 2, main text), while the dissociation constant of ATP, *K*_d_ (ATP), was taken from Rogne *et al*. and considered as not significantly impacted by urea.^13^ Explicit substrate binding and dissociation rates were treated as free parameters, but were required to maintain the experimental *K*_d_ values; AMP can bind to both the LID-open and -closed state; accordingly, *K*_d_ (AMP) is given by

$$K_{d}\left( AMP \right)=\frac{k_{\mathrm{off}}\left( AMP \right)}{k_{on}\left( AMP \right)},$$

(10)

with *k*_on_ (AMP) the binding rate constant for AMP and *k*_off_ (AMP) the dissociation rate. As ATP can bind and unbind only in the open state, we took into account the conformational equilibrium in the bound (*K*_C,ET_) and unbound (*K*_C,E_) species for the dissociation constant of ATP, *K*_d_ (ATP),

$$K_{d}\left( ATP \right)=\frac{k_{off}\left( ATP \right)\cdot\left( K_{C,E}+1 \right)}{k_{on}\left( ATP \right)\cdot\left( K_{C,ET}+1 \right)},$$

(11)

with *k*_on_ (ATP) the binding rate constant for ATP and *k*_off_ (ATP) as the dissociation rate. Lower limit for substrate dissociation rates of *k*_off_ (ATP) ≥ 544 s^‑1^ and *k*_off_ (AMP) ≥ 6340 s^‑1^, provided by Fry and coworkers,^15, 16^ were used in the fitting procedure.

Other free parameters were the rate of the actual phosphotransfer step (*k*_cat_) and the rate of the correct positioning of substrates in the "ATP first" path $k_{r}^{T-path}$ and the "AMP first" path $k_{r}^{M-path}$. These parameters were considered as unaffected by urea. While it is possible that urea also affects *k*_r_ and *k*_cat_, the impact is likely minimal as enzymatic velocity is accurately described with these parameters shared across different urea concentrations. All free parameters were optimized globally to match the enzymatic turnover as a function of either AMP or ATP concentration. In this procedure, we formulated a set of differential equations that describe the kinetics in Supporting Note 1. The concentration of any involved species was calculated by integrating these differential equations, which was done numerically using MATLAB's ordinary differential equation solver ode15s.^17^ The change of the substrate concentration over time represents the turnover *v_0_*. This calculation was repeated for different initial concentrations of urea and the substrates. Finally, we used a χ^2^-minimization of *v_0_* versus the substrate concentration to optimize the free parameters. The goodness of the fit in each case was judged based on the reduced χ^2^ values.

As expected, the optimized parameters for binding rate constants of *k*_on_ (ATP)  = 2.8·10^7^ M^-1^ s^‑1^ and *k*_on_ (AMP) = 1.9·10^7^ M^-1^ s^‑1^ are 1-2 orders of magnitude smaller than the diffusion limit, which was evaluated from the well-known Smoluchowski equation,

$$k_{D_{0}}=4\pi DR,$$

(12)

where *D* is the relative translational diffusion constant and *R* is the contact distance between the centers of the two molecules.

In the case of the mutant proteins L107I and F86W, the binding rate constants of the WT were provided as inputs, assuming that the encounter rate between substrate and enzyme is unaffected by the mutations. The dissociation rates were treated as free parameters, as the mutations likely affect the interactions between the protein and the bound substrate, in particular for F86W with the mutation in the AMP binding site.^18^ This means that for the mutant proteins the *K*_d_ values were not constrained. For F86W, which does not show substrate inhibition, a simplified model containing only the "ATP first" path was used. Our single-molecule experiments did not detect statistically significant differences (P >0.05, Student's t-test) in the protein dynamics between E and EM, as well as between ET and EMT_i_, suggesting that the "AMP first" path is only weakly populated. It is plausible that the mutation of F86 prevents the formation of the inactive EMT_i_ orientation of the nucleotides.

To estimate the confidence intervals of the fitted parameters, we monitored the increase in $\chi_{\mathrm{red}}^{2}$ ($\chi^{2}$ per degree of freedom) upon the perturbation of each parameter. For this, we used the optimized values from the χ^2^-minimization (Table 1, main text) and fixed the tested parameter at different values surrounding the optimal one. Then, the remaining parameters were optimized under this constraint. Afterwards, we calculated the difference in $\chi_{\mathrm{red}}^{2}$ with and without the constraint. A sharp increase in Δ$\chi_{\mathrm{red}}^{2}$ indicates that the goodness-of-fit is highly sensitive to the tested parameter. To distinguish between urea's effect on conformational dynamics and substrate affinity, we tested computationally how the enzymatic velocity would change when only one of the parameters is affected by urea at a concentration of 0.8 M while preserving all other parameters as determined at 0 M urea. Finally, to evaluate how enzymatic velocity depends on the actual opening and closing rates, we scaled the experimentally derived closing and opening rates of the ATP-bound species (ET, ETM_i_, EMT_i_, ETM) by a factor between 0.01 and 100. The conformational dynamics of species that are not bound to ATP (E, EM) were not altered as they are hardly affected by urea (Fig. 3c, main text).

**Supporting Figures**

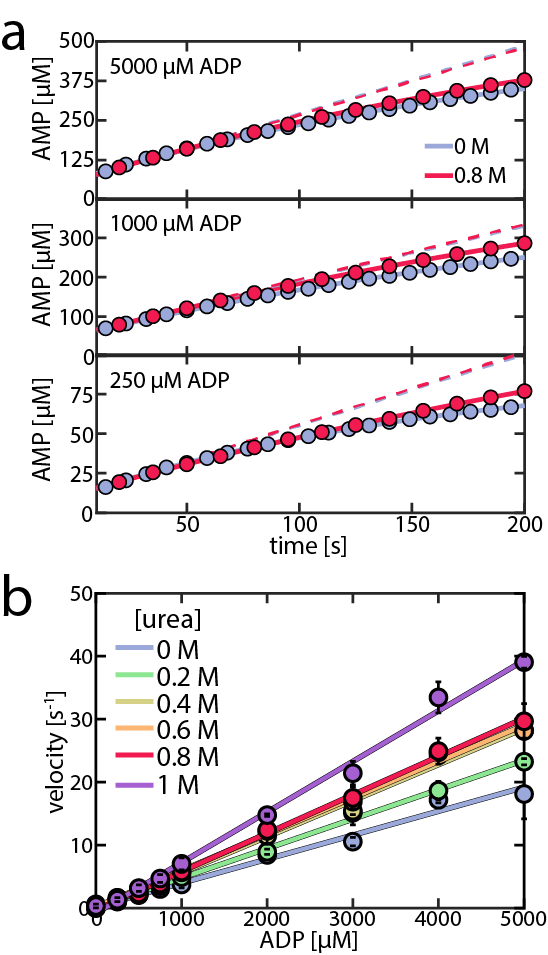

**Figure S1: Effect of urea on the AMP inhibition of the backward reaction**. a) Shown is the AMP concentration following the start of the reaction by injection of Mg^2+^ for representative initial ADP concentrations of 5000 µM (top), 1000 µM (center) and 250 µM (bottom) and for urea concentrations of 0 M (blue) and 0.8 M (red). The curves at 0.8 M urea represent the data shown in Fig. 1d in the main text. The solid curves represent a fit according to Eq. (1) (main text). The dashed lines represent the extrapolated product formation without product inhibition. While the initial velocities *v_0_* are not significantly affected by urea (Fig. 1d, main text), product inhibition by AMP is weaker in the presence of urea. b) Enzymatic velocity of WT AK as a function of ADP concentration in the presence of 10 mM AMP. For the backward reaction, AMP is a competitive inhibitor for the ADP/AMP binding site, resulting in a strong reduction of turnover and an apparent linear dependence on the ADP concentration. Urea (blue to red curves) gradually alleviates the inhibition by AMP. Error bars indicate the standard error of the mean of at least 2 measurements.

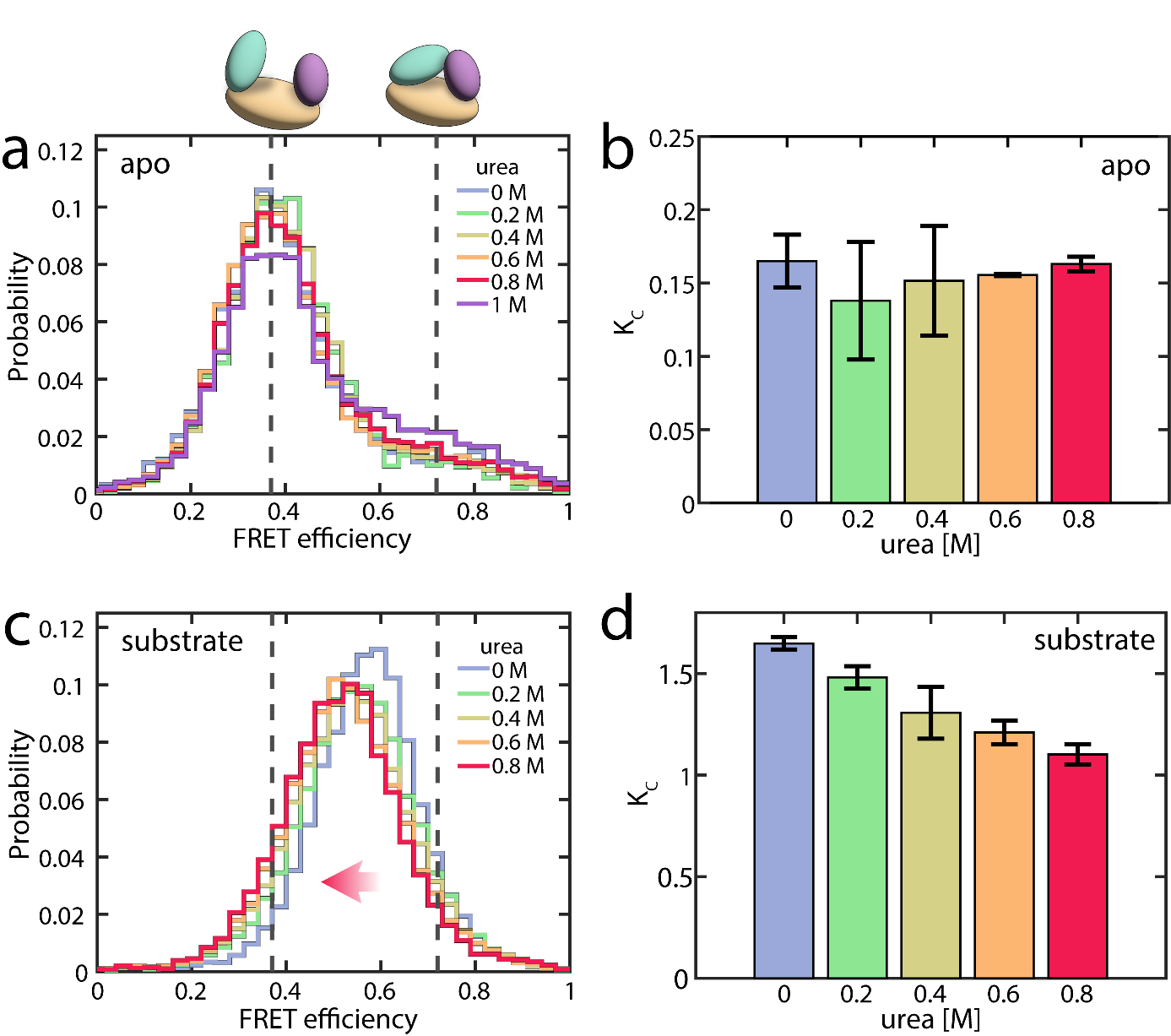

**Figure S2: Effect of urea on AK's conformation without and with substrate.** a) FRET efficiency histograms of the apoprotein at a series of urea concentrations. The dashed grey lines indicate the FRET efficiency values of the open (0.37±0.01) and closed (0.72±0.01) states obtained from the H^2^MM analysis. The impact of urea on the FRET efficiency histogram up to a concentration of 0.8 M is minor. A more significant effect is seen at a concentration of 1 M urea (violet), likely indicating the onset of global unfolding. b) Change in the equilibrium coefficient *K_C_* ratio as determined by H^2^MM for the apoprotein. c) In the presence of 1 mM ATP, 5 mM AMP and 417 µM ADP, increasing urea concentrations shift the histogram towards lower FRET efficiencies. d). As in (b), in the presence of substrates. Error bars indicate the standard error of the mean of at least 2 repeated measurements.

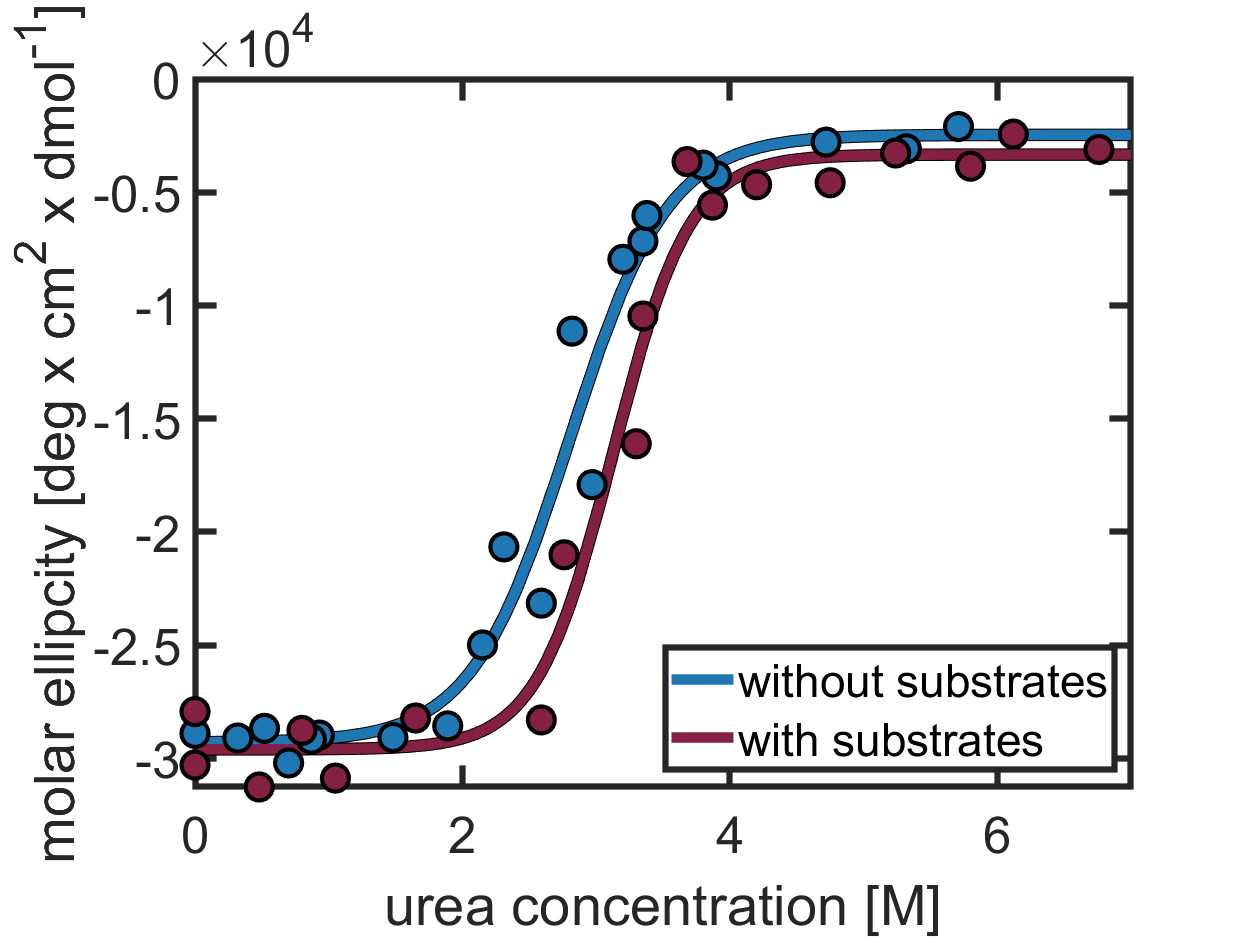

**Figure S3: Stabilities against urea-induced unfolding.** a) The unfolding of secondary structure elements in AK (AK C77S/V142C/A73C) was monitored by the CD signal at 222 nm in the absence (blue) or presence (red) of substrates. In the concentration range used for the enzymatic essays and smFRET experiments (0-0.8 M urea), the α-helical content of AK does not change, similarly as for the WT protein.^19, 20^ The solid lines indicate fits to a sigmoidal function with flat baselines( Eq. (8)). The resulting values for the Gibbs free energy of folding, ${\Delta G}_{\mathrm{fold}}^{0}$, the $m$-value for denaturant-dependent unfolding and the midpoint of the transition are given in Table S6. The binding of substrates increased the stability of AK, as observed before.^21^

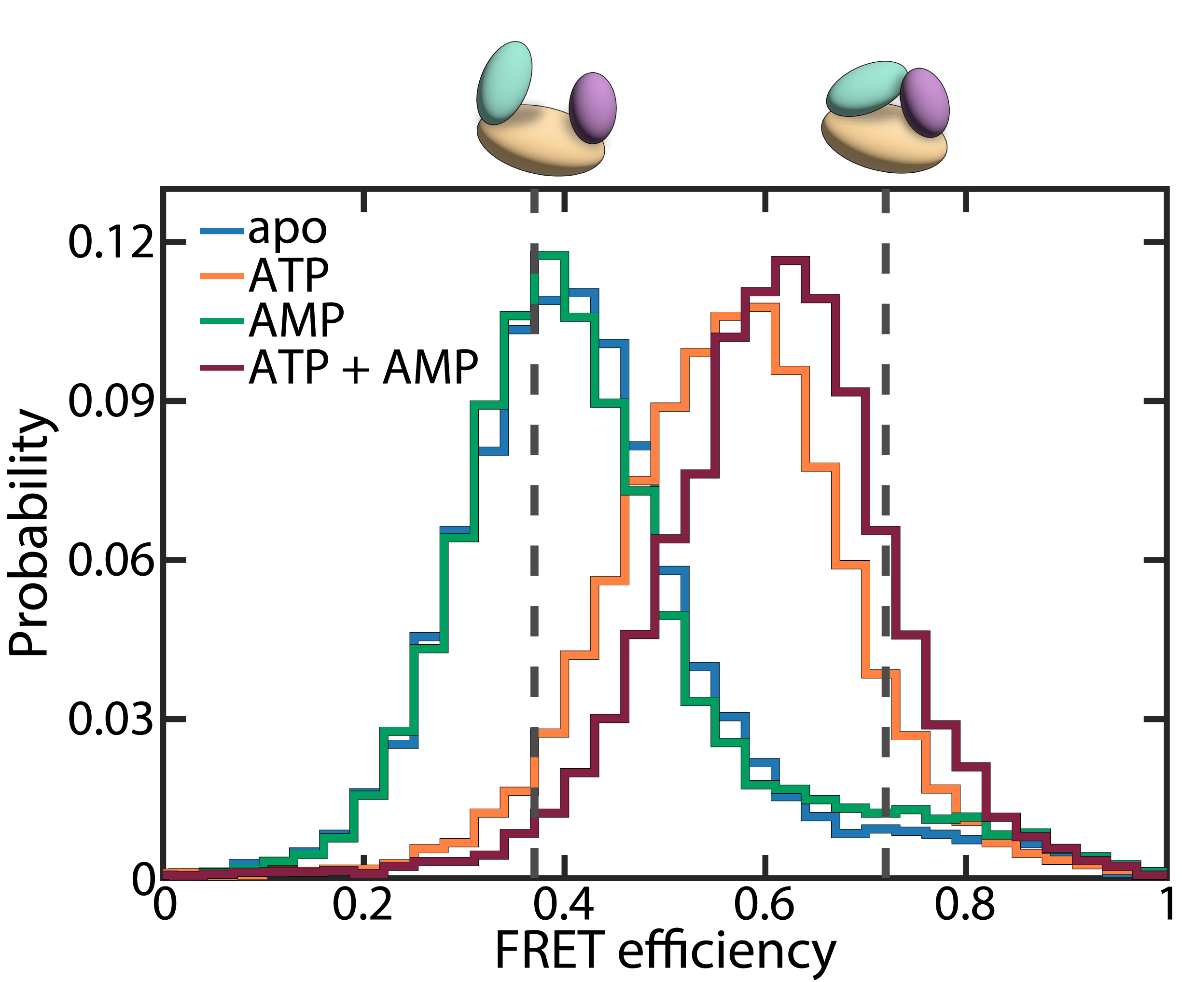

**Figure S4: Substrate-dependent domain closure.** The apoprotein (blue trace) mainly adopted an open conformation, occasionally exploring the closed state. AMP as a single substrate (5 mM, green) did not trigger significant closure of the LID domain. The binding of ATP (1 mM, orange) increased the population of the closed state. The additional presence of AMP (5 mM, red) shifted the histogram even more towards higher FRET efficiencies. The dashed grey lines indicate the FRET efficiencies of the open (0.37±0.01) and closed (0.72±0.01) state according to H^2^MM analysis.^1^

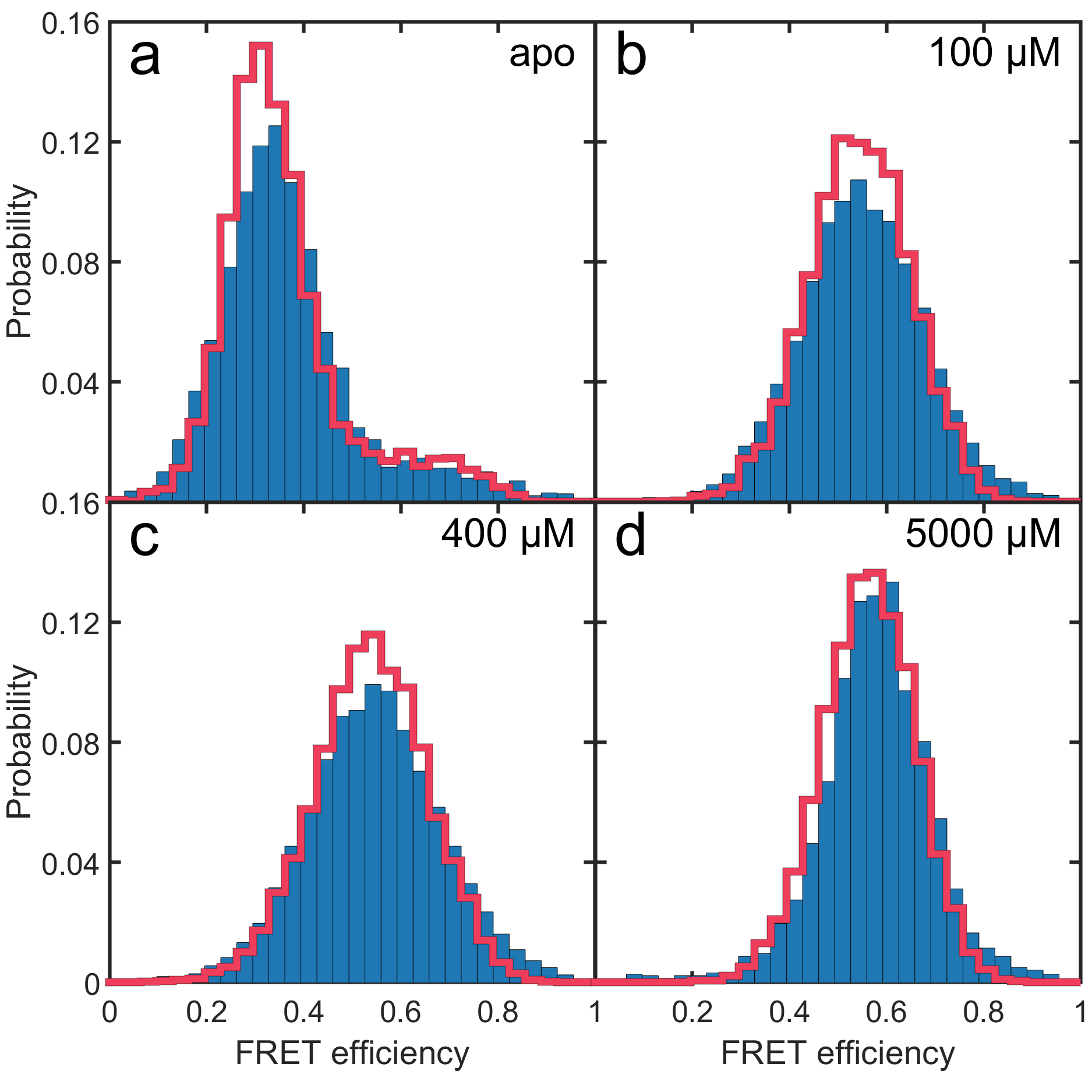

**Figure S5: Validating H^2^MM models using recoloring.** Representative experimental histograms are shown in blue for different AMP concentrations, as indicated in each panel. The recolored histograms are depicted as solid red lines and show good agreement between simulation and experiment. In b-d), the ATP concentration was fixed at 1 mM, and the ADP concentration was adjusted to guarantee equilibrium (Table S4).

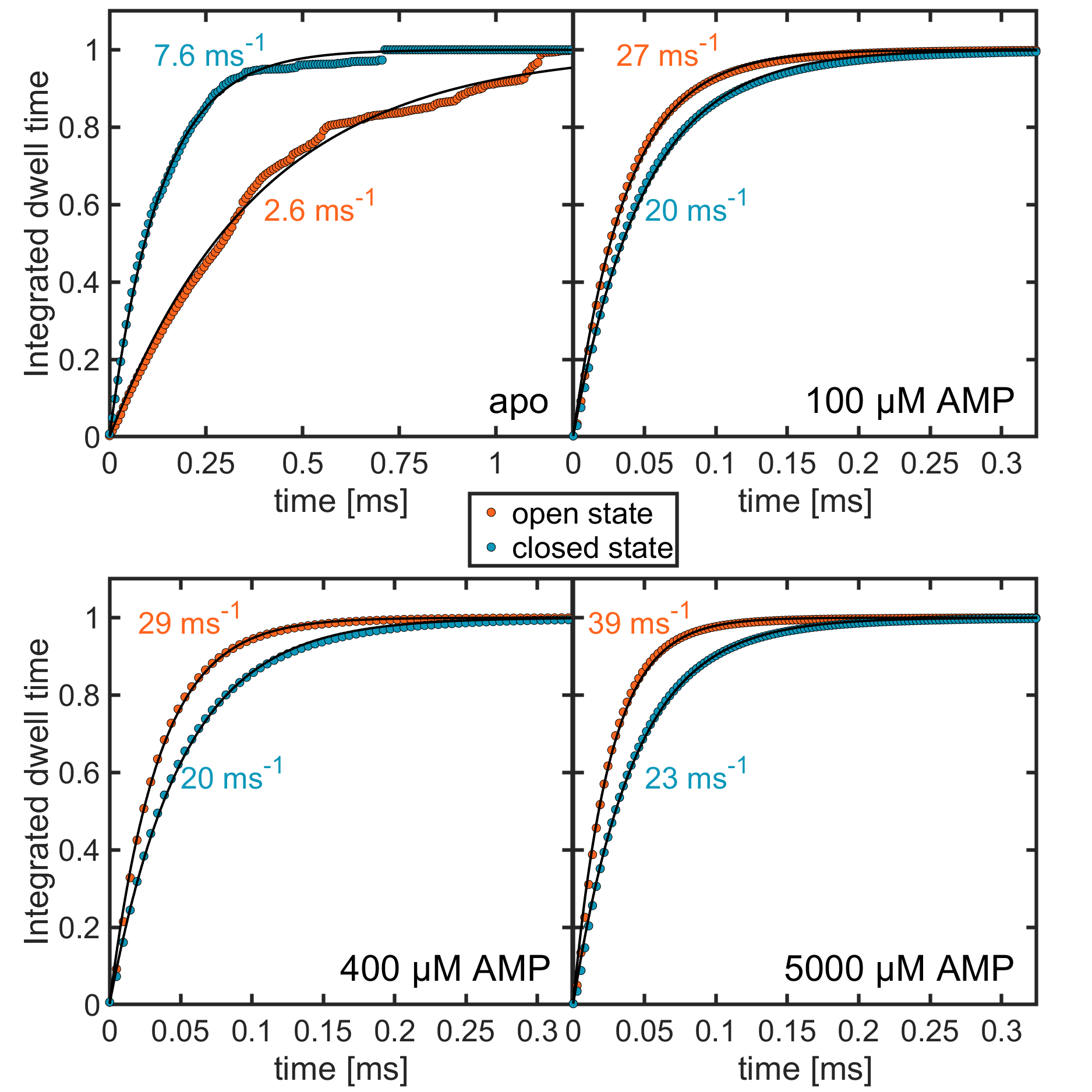

**Figure S6: Dwell-time analysis.** Integrated dwell-time distributions are shown for the open state (orange) and the closed state (cyan) at four different concentrations of AMP, as indicated within each panel. In b-d), the ATP concentration was fixed at 1 mM, with the ADP concentration adjusted to maintain equilibrium. Black lines represent fits to single-exponential functions. Both closing and opening rates extracted from the dwell time distributions compare favorably to the rates obtained by H^2^MM analysis (Table S3).

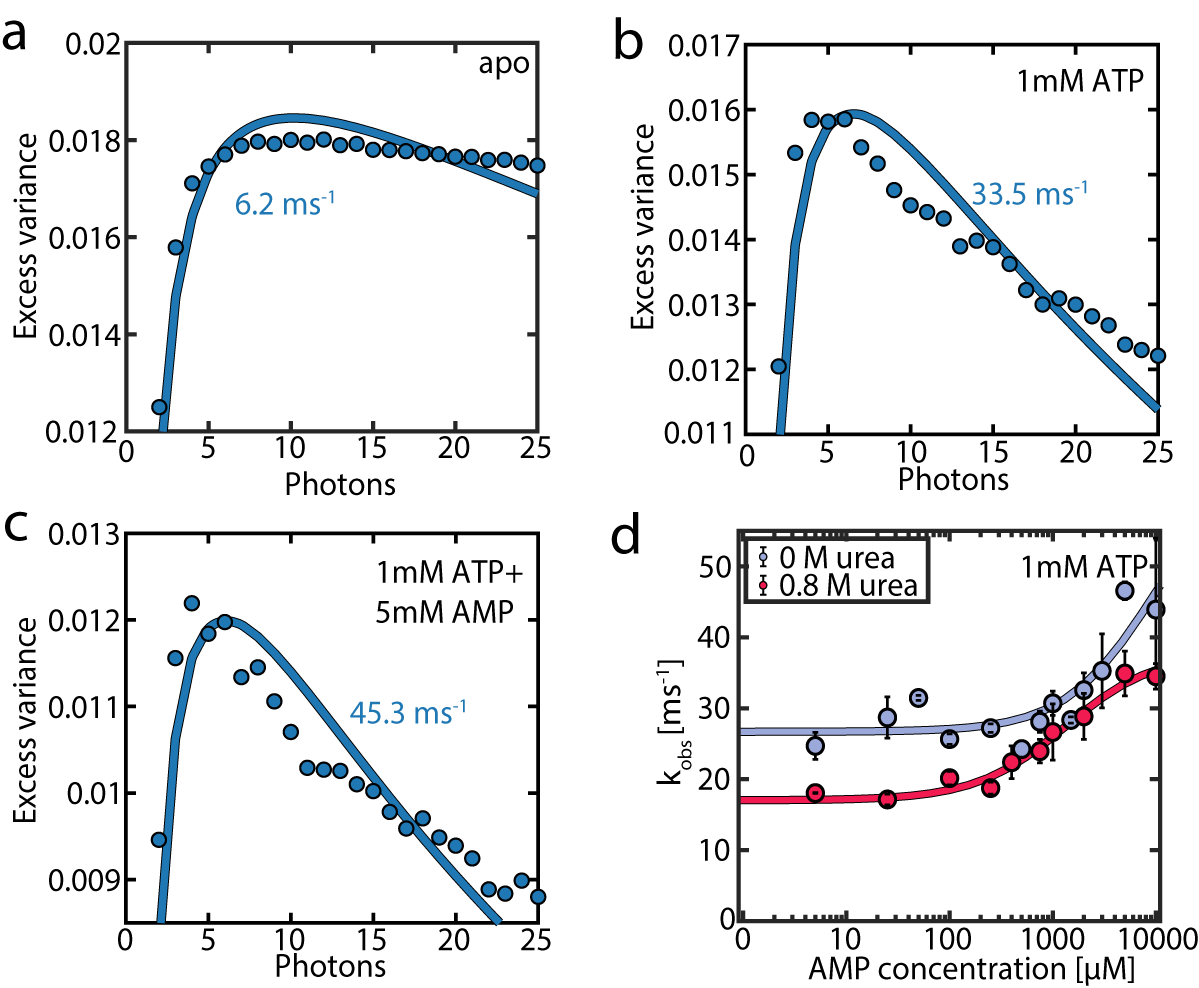

**Figure S7: Time-resolved burst variance analysis (**trBVA**)**. a-c) Traces of the excess variance with increasing numbers in the photon segments.^10^ Solid lines represent fits to a model with 2 conformational states. The fits have two fitting parameters, the amplitude <δ*ε*^2^> and the observed rate *k*_obs_ = *k*_o_ + *k*_c_. trBVA confirms the presence of protein dynamics on the µs-time scale, which are accelerated by adding substrates. d) The observed rate *k*_obs_ as a function of AMP concentration for a fixed concentration of ATP (1 mM). The data set used is the same data set used to generate Fig. 4 in the main text. The solid lines indicate a fit to a binding isotherm. In agreement with H^2^MM analysis, in the absence of AMP, *k*_obs_ is slowed down in the presence of urea (red curve). Likewise, the rates are accelerated by adding AMP, both in the presence and absence of urea. The observed rates reported in (d) are ~2 times slower than the sum *k*_o_ + *k*_c_ in the H^2^MM results (Fig. 4), a result that lies within the expected deviation from the ground truth for trBVA.^10^

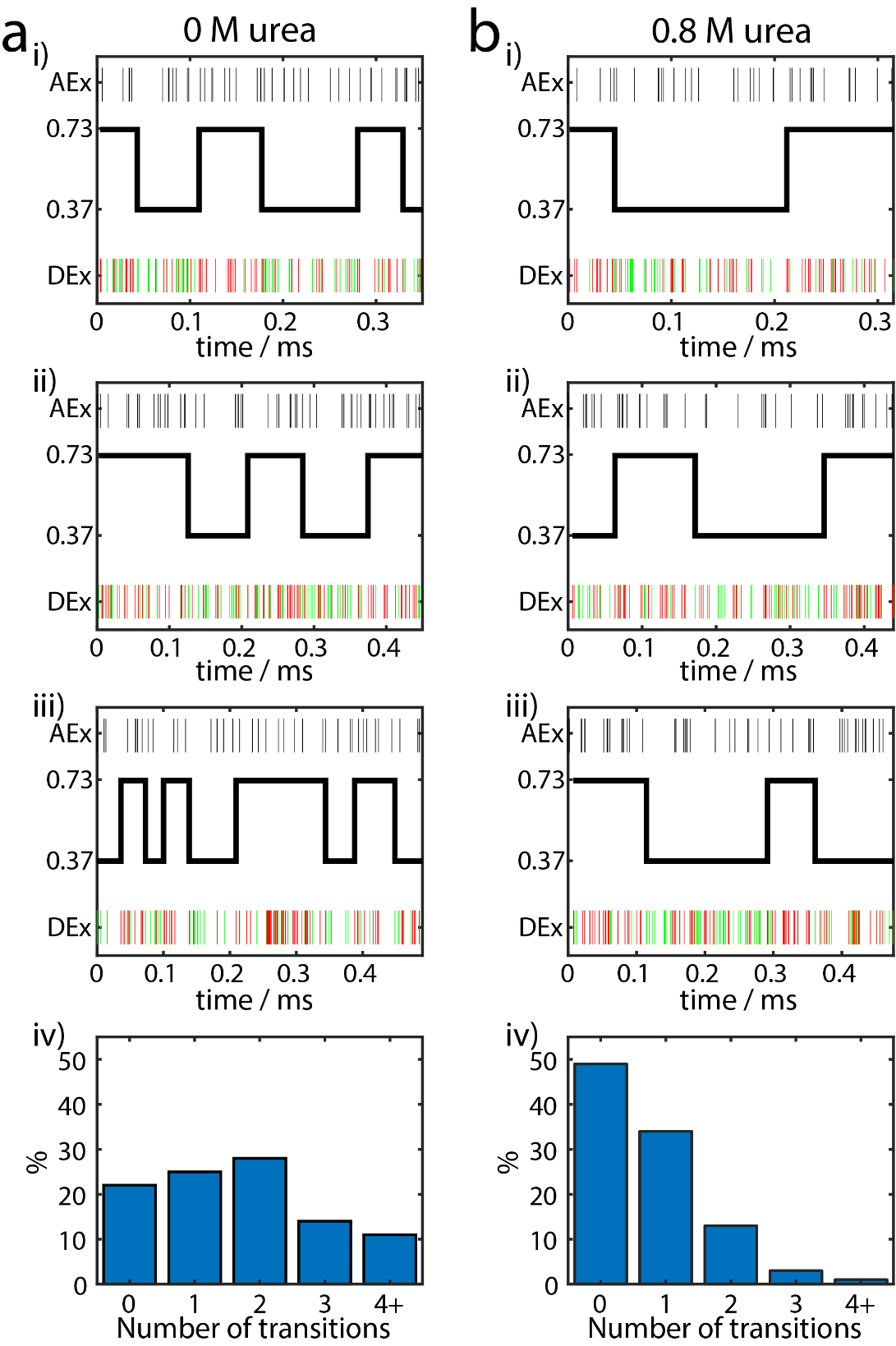

**Figure S8: Representative photon-by-photon trajectories and Viterbi assignments**. Single-molecule trajectories are shown for the WT protein at 1 mM ATP in (a) the absence of urea and (b) the presence of 0.8 M urea. For i-iii), the top panel shows the arrival time of photons after the acceptor pulse. The bottom panel shows the arrival time of photons after the donor pulse, in green for donor photons and red for acceptor photons. The black solid line depicts the most likely state sequence according to a Viterbi assignment.^8^ iv) depicts the distribution of the number of transitions identified by the Viterbi algorithm in each data set. In the presence of only ATP, urea slows down the transition rates (main text Fig. 4), leading to fewer transitions in (b).

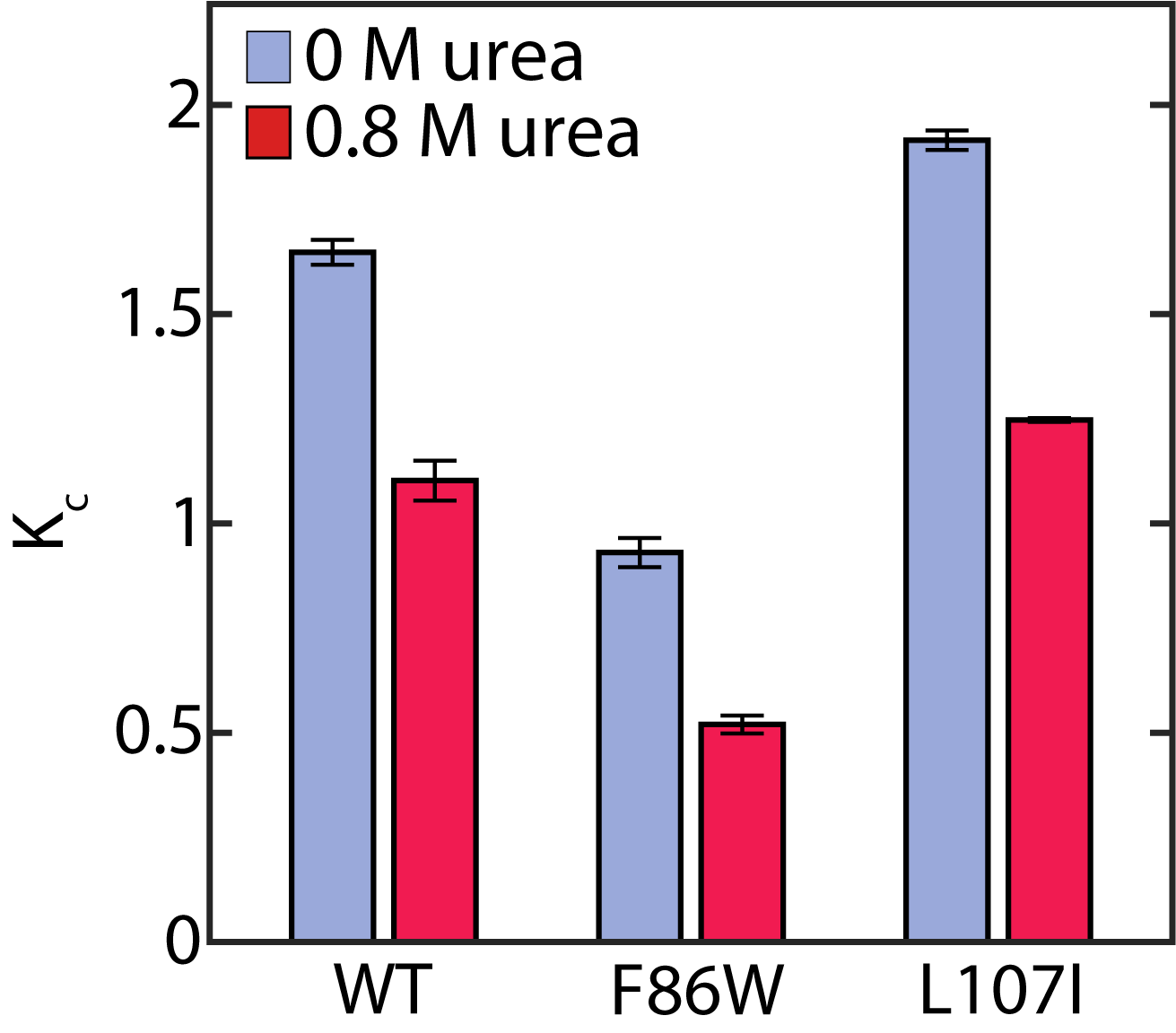

**Figure S9: The effect of urea on the occupancy of the closed state for the mutant proteins.** a) The equilibrium coefficient *K_C_* for the mutant proteins F86W and L107I at with 1 mM ATP, 5 mM AMP and 417 µM ADP. The non-inhibited F86W and the strongly inhibited L107I are both shifted towards the open state by urea. Also, L107I has a higher occupancy of the closed state than the WT, while for F86W *K*_C_ is lower.

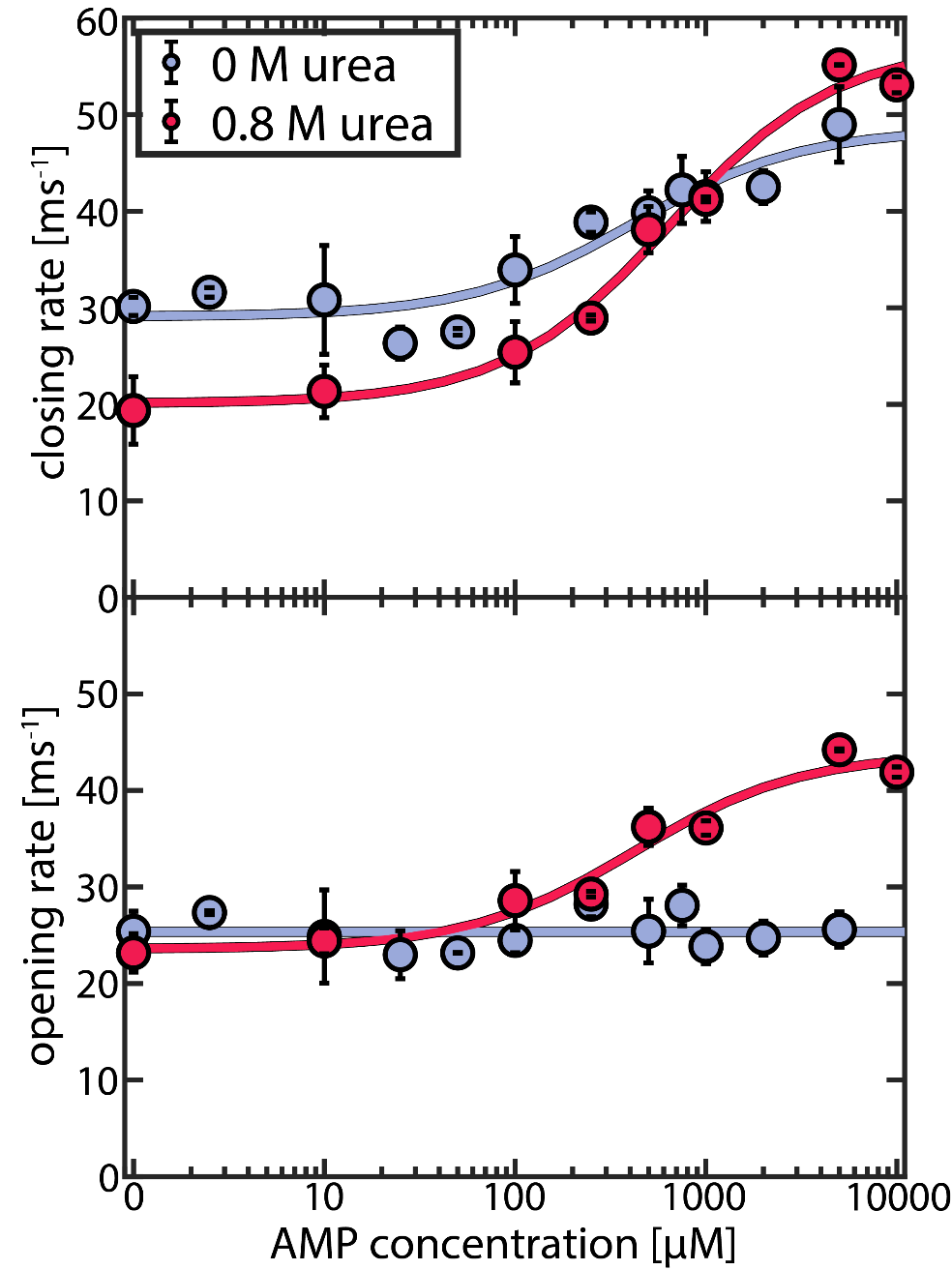

**Figure S10: AMP-dependent closing and opening rates for L107I.** Closing (upper panel) and opening (lower panel) rates for the L107I protein as a function of AMP concentration. Experiments were conducted at a fixed ATP concentration of 1 mM. Blue corresponds to the absence of urea, and red to the presence of 0.8 M urea. Solid lines indicate fits to a model described in "Supporting Note 2: Analysis of binding-state dependent opening and closing rates". The error bars indicate the standard error of the mean of at least 2 measurements.

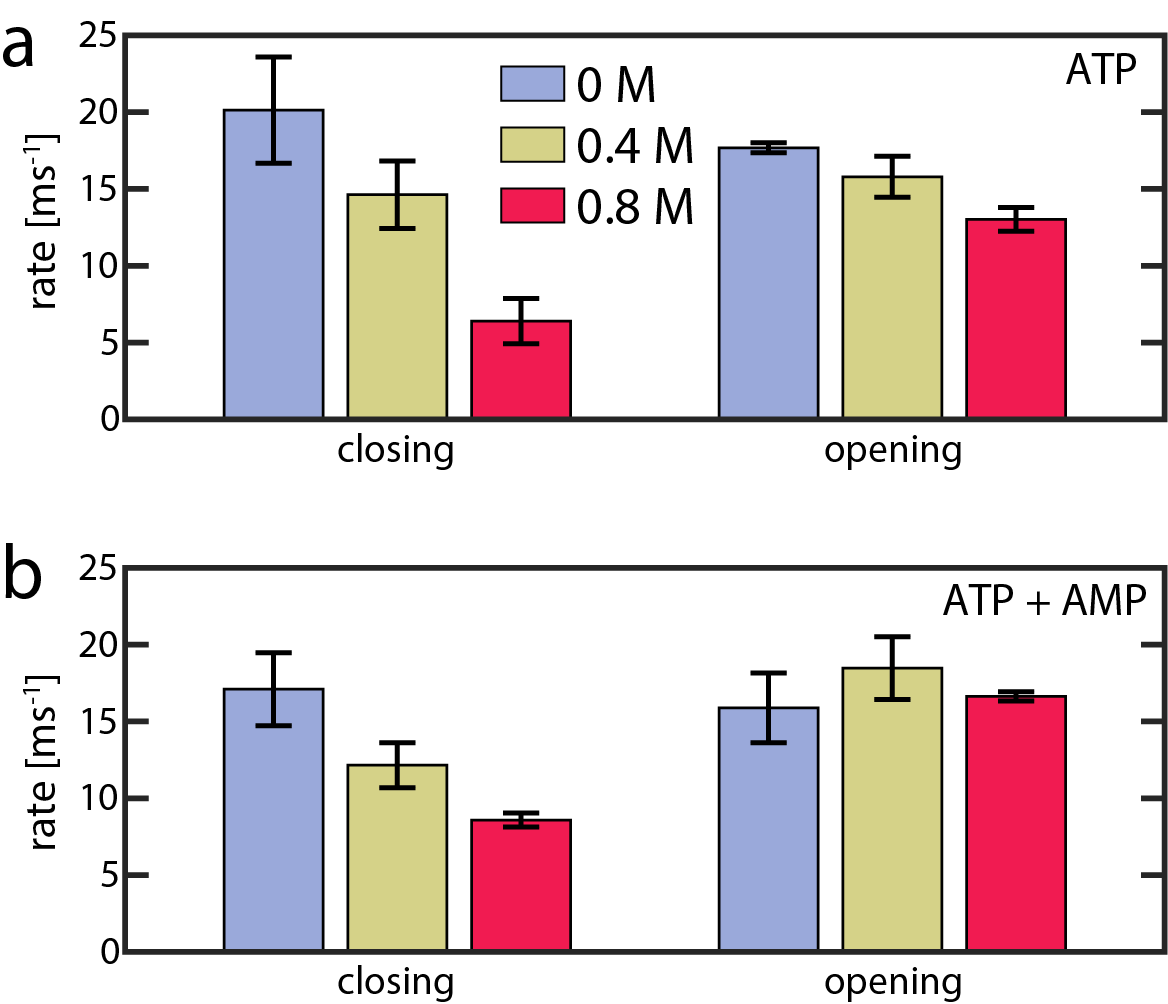

**Figure S11: Effect of urea on the closing and opening rates of F86W.** Closing (upper panel) and opening (lower panel) rates for the F86W protein as a function of the urea concentration, for a) 1000 μM ATP and b) 1000 μM ATP, 5000 μM AMP and 417 µM ADP. The error bars indicate the standard error of the mean of at least 2 measurements. Blue corresponds to the absence of urea, yellow to 0.4 M urea, and red to 0.8 M urea. For F86W, urea had a more pronounced impact on domain closing than opening, similar to both the WT (Fig. 4, main text) and L107I (Fig. S10) in the absence of inhibiting concentrations of AMP. Furthermore, comparing the values between (a) and (b) indicated that the addition of AMP to the ATP-bound protein did not cause statistically significant changes (P >0.05, Student's t-test), unlike in the other proteins in the presence of high concentrations of AMP (Fig. 4 main text, Fig. S10).^1^ This suggested that the "AMP first" pathway is only very weakly populated or not at all in F86W.

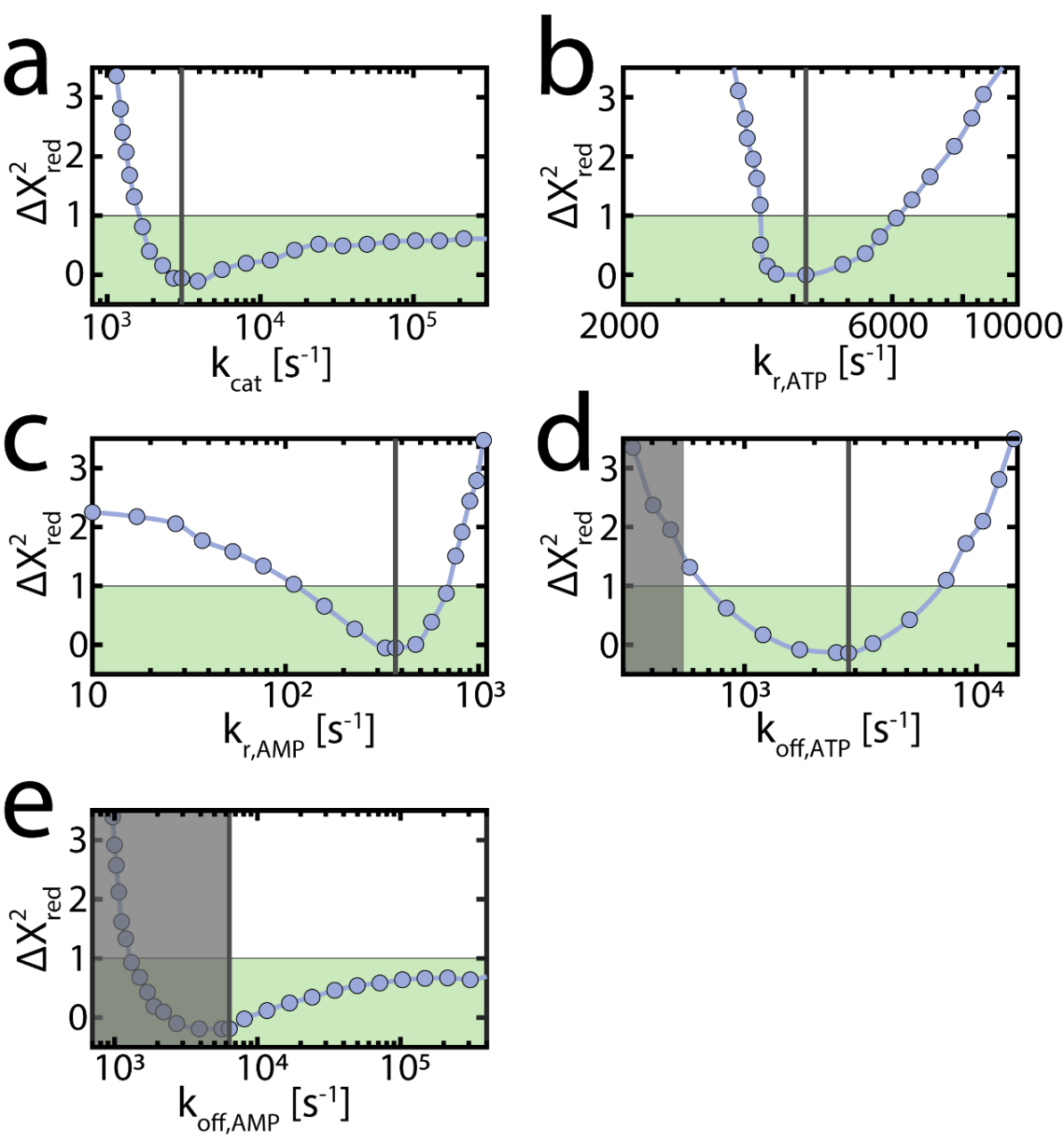

**Figure S12: Confidence limits of parameters describing the enzymatic activity of WT AK.** To estimate the parameters describing the enzymatic activity of AK, we used a *χ*^2^-minimization. These parameters are a) *k*_cat_, b) *k*_r,ATP_ and c) *k*_r,AMP_. We additionally fitted in (d) and (e) the explicit values for the dissociation rates *k*_off,ATP_ and *k*_off,AMP_, with a lower limit taken from NMR experiments indicated by the grey box.^15, 16^ Nucleotide binding rates were derived from the dissociation rates together with the measured *K*_D_ values. To estimate the confidence intervals of the fitted parameters, we monitored the increase in $\chi^{2}$ per degree of freedom, $\chi_{\mathrm{red}}^{2}$, upon the perturbation of each parameter. For this, we used the optimized values from the χ^2^ minimization (indicated by the black lines) and fixed the tested parameter at different values surrounding the optimal one. Then, the remaining parameters were optimized under this constraint, and we calculated the difference in $\chi_{\mathrm{red}}^{2}$ with and without the constraint, ${\Delta\chi}_{red}^{2}$, indicated by the blue dots. A ${\Delta\chi}_{red}^{2}$ value of 1 represents a 68.3 % confidence interval^22^. For example, a sharp increase in ${\Delta\chi}_{red}^{2}$ was observed when *k*_cat_ dropped below 1.5·10^3^ s^-1^ (panel a). In contrast, the upper limit was less reliable, as a fast catalysis could be compensated to some extent by other parameters, in particular a faster rate for *k*_off,AMP_. Notably, this analysis did not take into account physical reasonableness of the parameters; a very high rate for *k*_off,AMP_ means that *k*_on,AMP_ would exceed the diffusion limit in order to maintain experimental *K*_D_ values, which is unlikely.

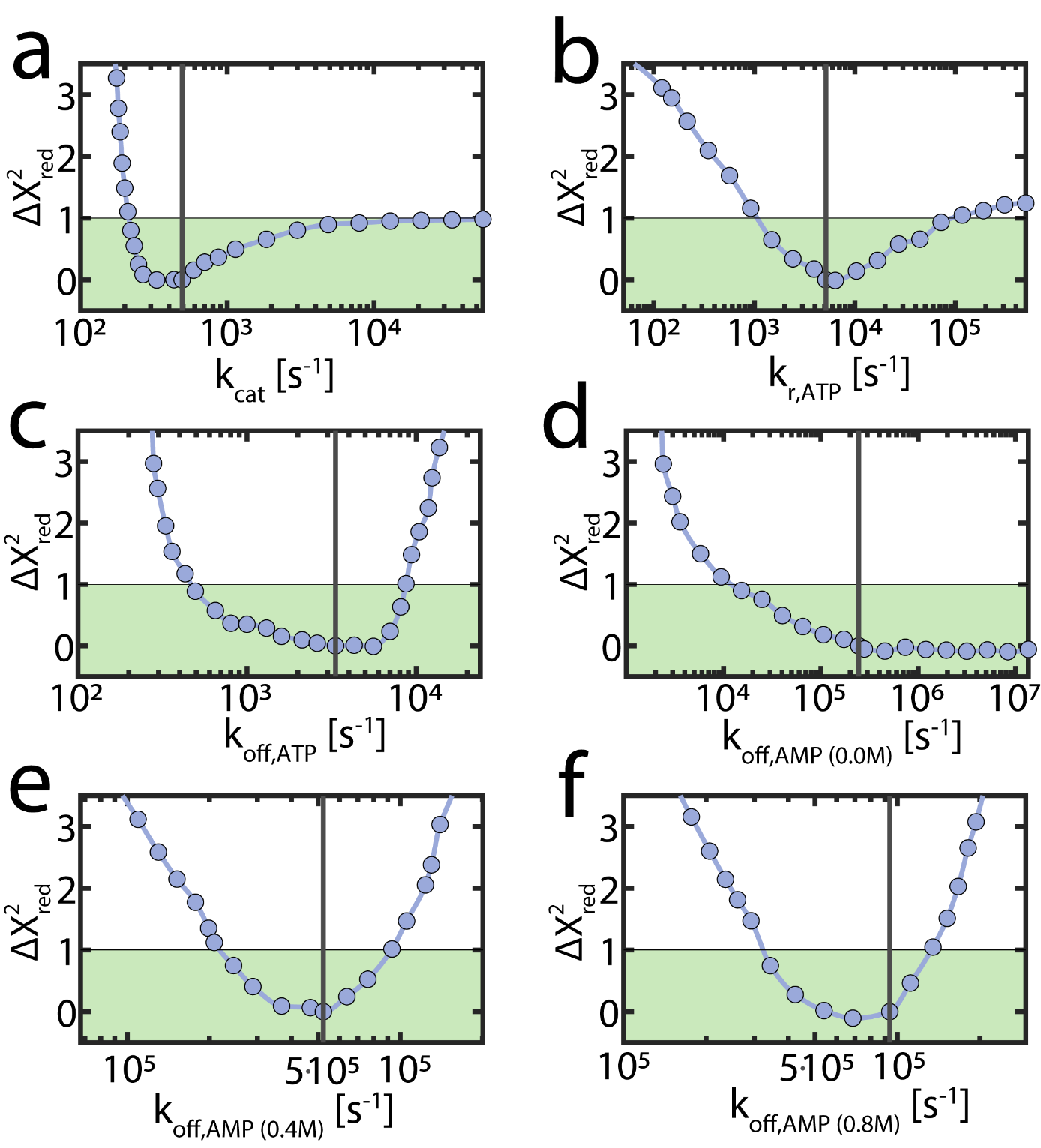

**Figure S13: Confidence limits for the enzymatic parameters of the non-inhibited mutant F86W.** As in Fig. S12, the change in $\chi_{red}^{2}$ upon perturbation of the optimal values (black lines) was monitored. These parameters are a) *k*_cat_, b) *k*_r, ATP_, c) *k*_off, ATP_ and d-f) *k*_off, AMP_. In contrast to Fig. S12, the dissociation rates were fitted without applying a restriction on *K*_D_, to accommodate for potential changes in nucleotide affinity introduced by the mutation. F86 is located close to the AMP-binding site, causing a severe drop in affinity for AMP, while the affinity for ATP is weakly affected, in agreement with results of Liang *et al*.^18^ For the non-inhibited F86W, the enzymatic velocity could be fitted with a simplified model containing only the "ATP first" pathway. A ${\Delta\chi}_{red}^{2}$ value of 1 represents a 68.3 % confidence interval, which is indicated by the green box.

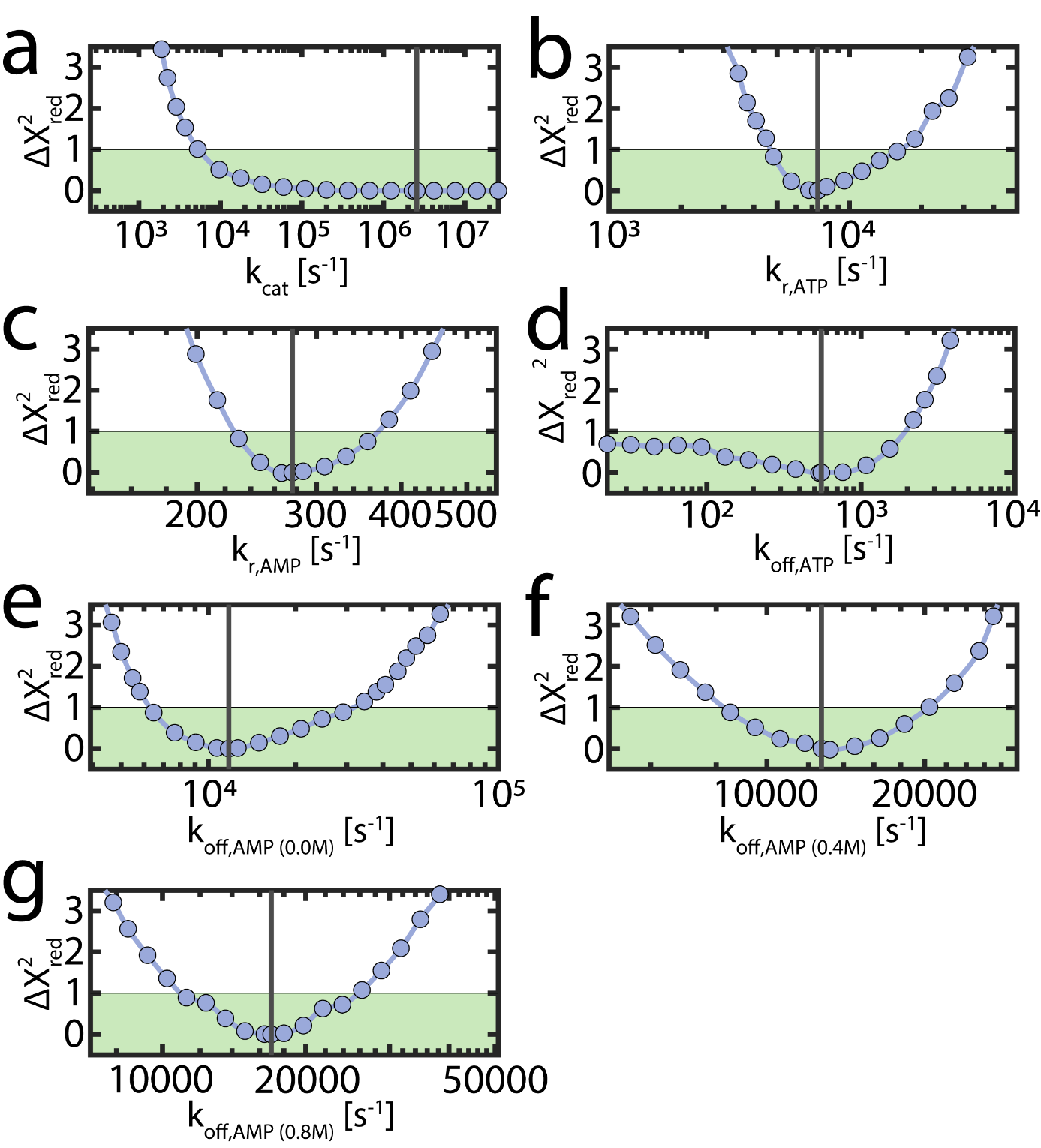

**Figure S14: Confidence limits for the enzymatic parameters of the strongly inhibited mutant L107I.** As in Fig. S12, the change in $\chi_{red}^{2}$ upon perturbation of the optimal values (black lines) was monitored. These parameters are a) *k*_cat_, b) *k*_r,ATP_ , c) *k*_r,AMP_, d) *k*_off, ATP_ and e-g) *k*_off, AMP_. The dissociation rates were fitted without applying a restriction on *K*_D_, to accommodate for potential changes in nucleotide affinity introduced by the mutation. A ${\Delta\chi}_{red}^{2}$ value of 1 represents a 68.3 % confidence interval and is indicated by the green box.

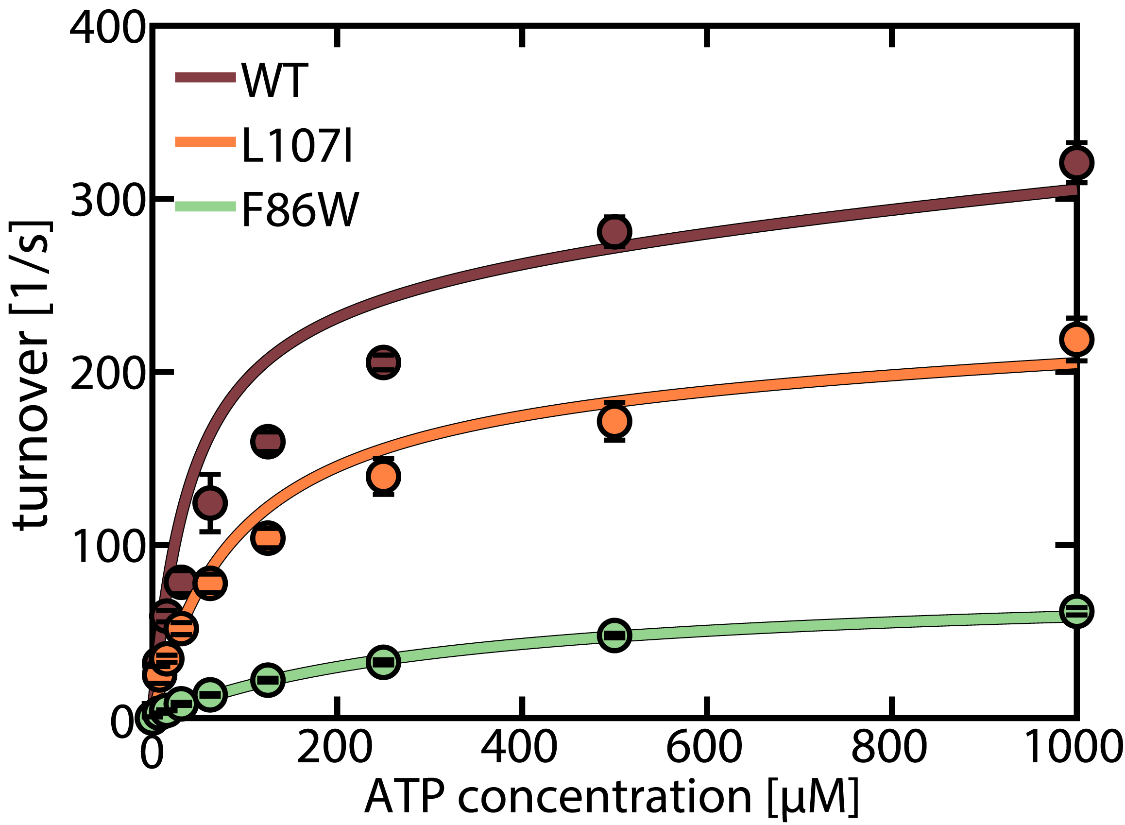

**Figure S15. ATP-dependent activity for AK mutants.** Enzymatic velocity as a function of ATP concentration for a fixed concentration of 1000 μM AMP and in the absence of urea. Shown are the WT protein (brown), L107I (orange) and F86W (green). The straight lines represent fits to the model described in "Supporting Note 1: Model for the substrate inhibition by AMP". The fit parameters are shared with the AMP-dependent activity (Fig. 1a-c main text) and given in Table 1 (main text).

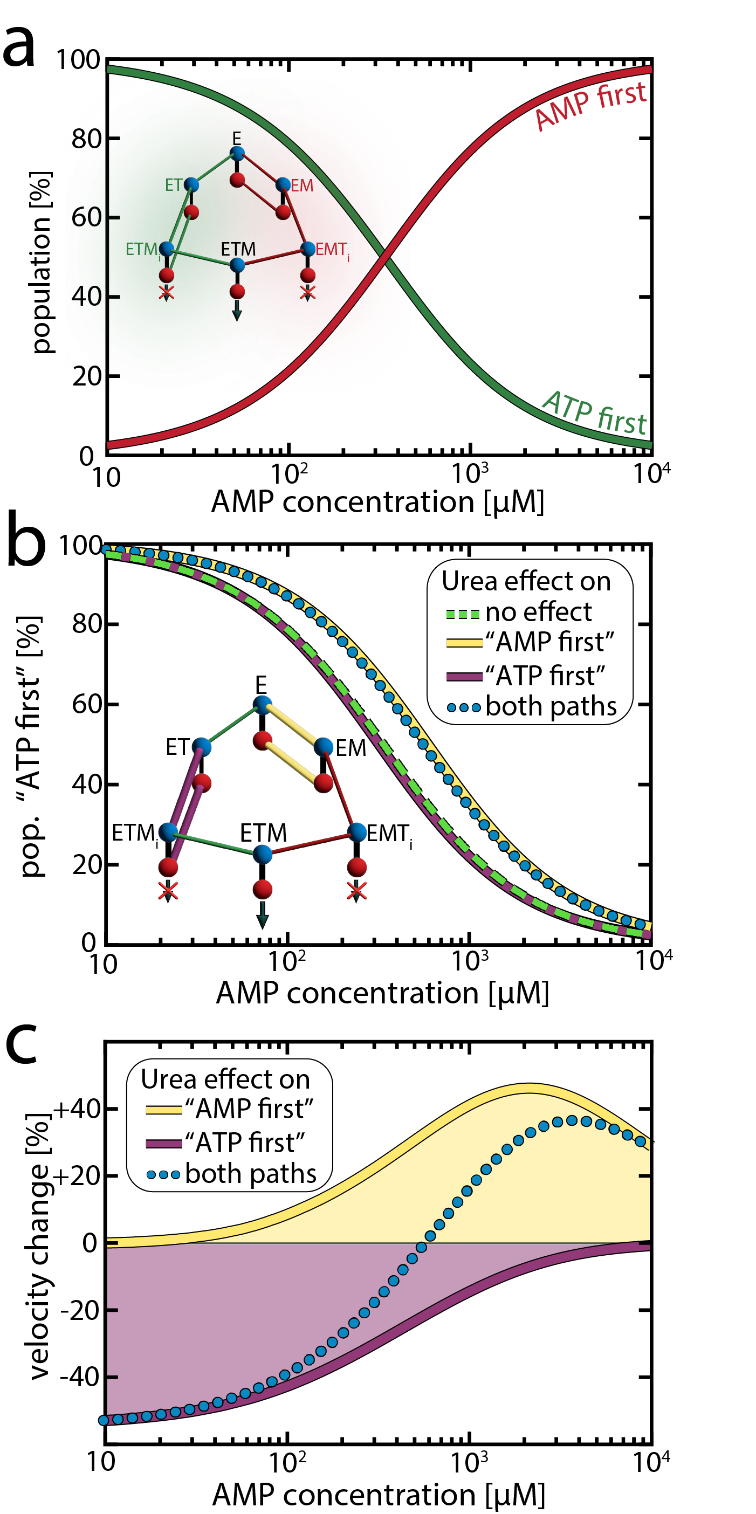

**Figure S16. Population of the competing "ATP first" and "AMP first" pathways.** To understand how the urea-induced reduction in AMP affinity can increase velocity, it is helpful to understand how the enzyme population is distributed between the two pathways.

a) The population of the two pathways as a function of the AMP concentration at 0 M urea, simulated according to our model for a fixed ATP concentration of 1 mM. The "ATP first" path in green contains the ET and ETM_i_ states and is dominating when the AMP concentration is small. The "AMP first" path in red contains the EM and EMT_i_ states and is becoming more populated as the AMP concentration increases. Both pathways are equally populated at 300-400 μM AMP, where the highest enzymatic velocity was observed (main text Fig. 1a). Further increasing the AMP concentration reduces the overall velocity, as the productivity loss in the "ATP first" path is more significant than the gain in the "AMP first" path due to the low $k_{r}^{M-path}$.

b) Change in the population of the "ATP first" path due to presence of urea. As in (a), we simulated the distribution of populations between the pathways. The dashed green curve depicts the population of the “ATP first” path in the absence of urea. To obtain the dotted blue line, we altered AMP affinity according to experimental observations, while preserving all other parameters, including those describing conformational dynamics, as determined at 0 M urea. The purple and yellow lines illustrate the impact on the population of the “ATP first” path when urea affects AMP affinity in only one of the two pathways. Although AMP binding is required in both pathways, decreasing its affinity had distinct effects in the two pathways. Increasing *K*_d_ (AMP) in the first step of "AMP first" path (highlighted in yellow in the inset) while preserving *K*_d_ (AMP) in the opposing path increased the likelihood of reaching the more productive "ATP first" path. In contrast, reducing the AMP affinity in the second step of the more productive "ATP first" path (highlighted in purple in the inset) hardly affected the distribution between the pathways.

c) Outcome of the urea-induced changes in the distribution between the pathways on enzymatic velocity. As shown in (b), reducing AMP affinity affected the distribution between the pathways differently depending on which pathway is affected. We simulated how these changes affect the overall enzymatic velocity. The yellow curve depicts the velocity change when urea affected only the affinity within the "AMP first" path (yellow), with AMP affinity in the opposing "ATP first" path preserved. The outcomes at low AMP concentrations were negligible, as the "AMP first" path was scarcely populated. However, at intermediate concentrations, the overall velocity increased due to a higher population of the "ATP first" pathway (b). At very high AMP concentrations, the impact of urea was weak, as AMP binding was much faster than dissociation. In contrast, reducing AMP affinity within the "ATP first" path (purple) had a purely detrimental effect on velocity, as it reduced the flux through the productive pathway. The effect was strongest at low AMP levels, where this pathway was most populated, and lost significance as the "AMP first" path was more populated. The dotted blue line shows the velocity change when the affinity is reduced in both pathways.

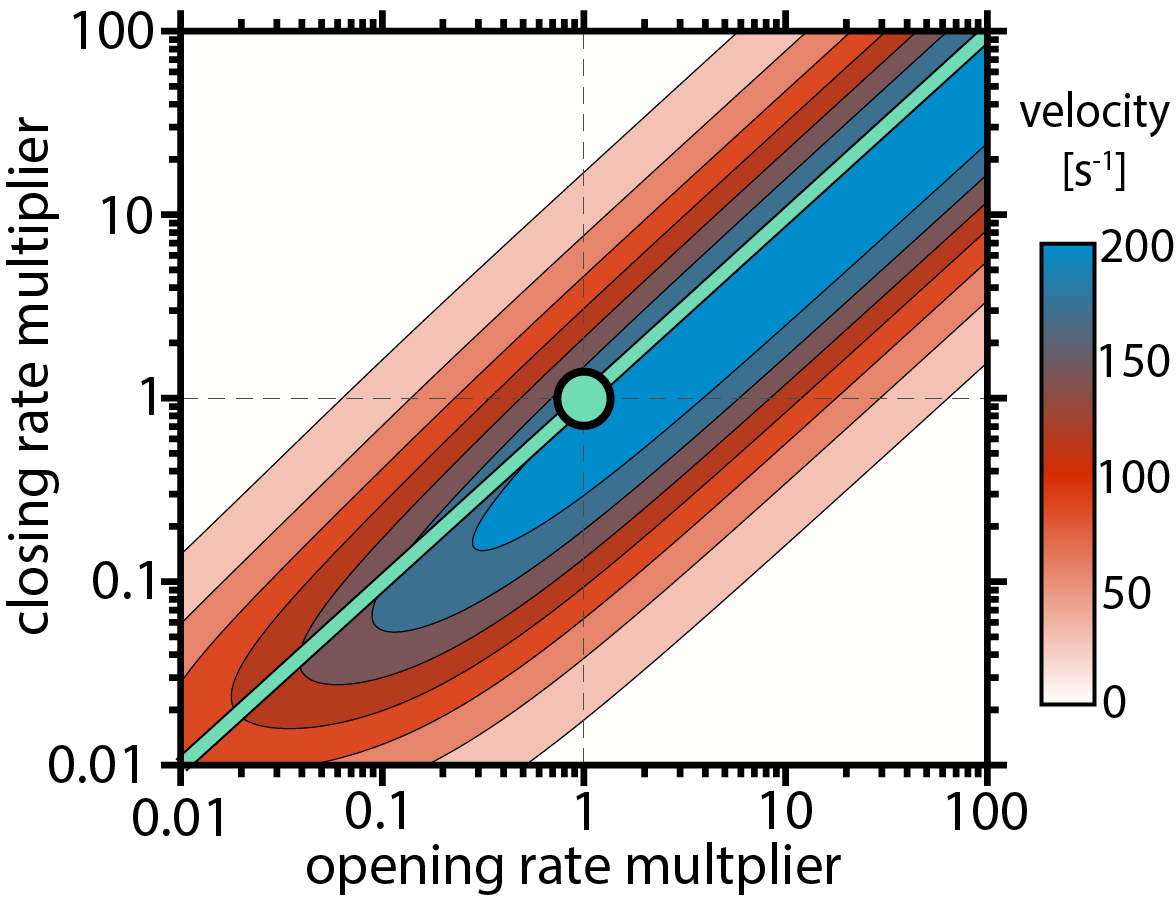

**Figure S17. Enzymatic velocity as a function of the opening and closing rate at physiological concentrations of nucleotides.** We simulated how the velocity changes when open and closing rates of the ATP-bound species (ET, ETM_i_, EMT_i_, ETM) are altered. The experimentally derived *k*_o_ and *k*_c_ were scaled by a factor between 0.01 and 100. Substrate concentrations of 0.3 mM AMP and 5 mM ATP reflected the physiological concentrations in *E.coli*.^23^ The optimal open/closed ratio obtained in the calculation was close to the experimental *K*_C_ value (turquoise circle), suggesting the enzyme has evolved to optimize *K*_C_ for maximal turnover under physiological substrate concentrations.

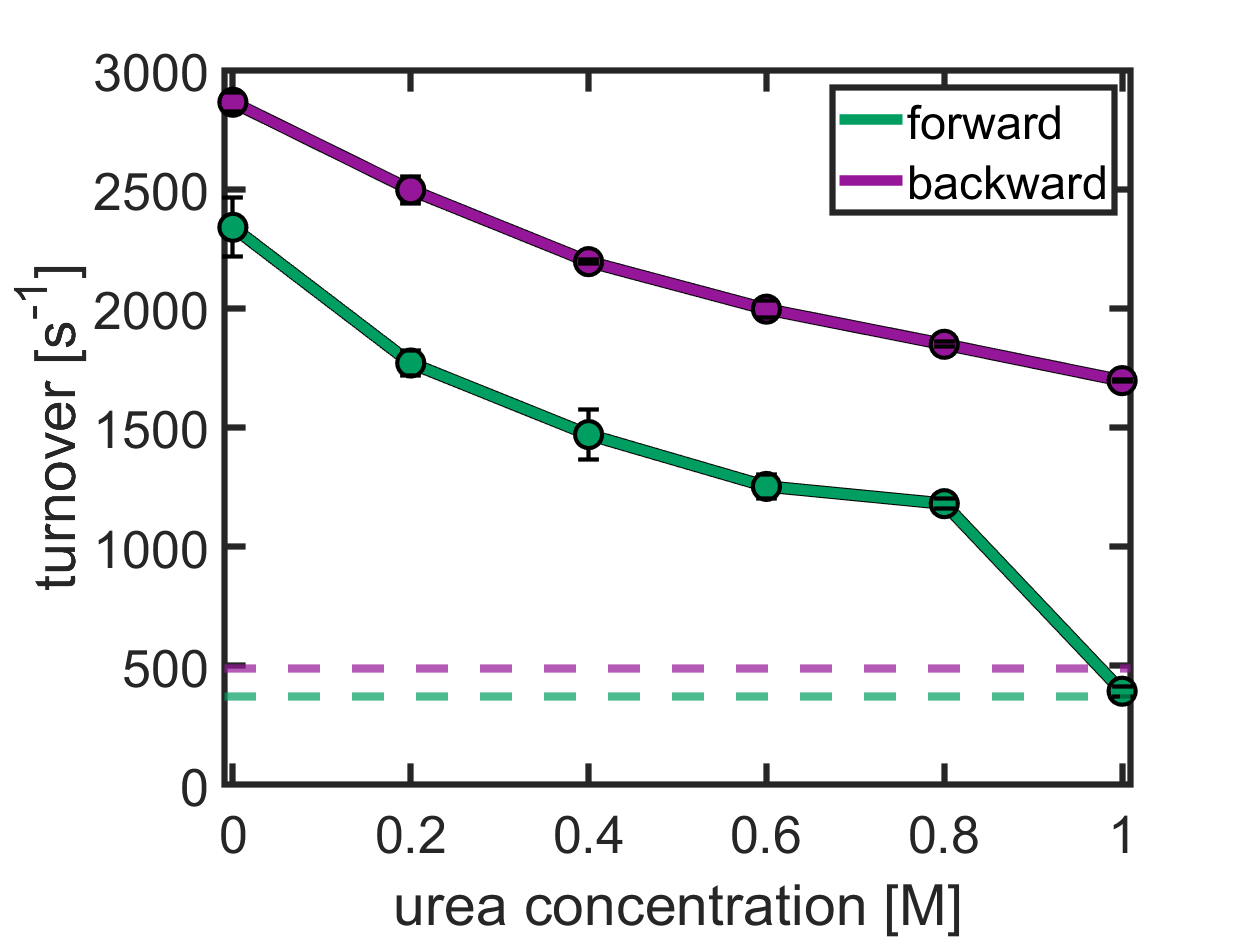

**Figure S18: Activity of coupled enzymatic systems.** A sufficient activity of the coupled enzymatic systems in the presence of urea must be guaranteed for a reliable assessment of AK's enzymatic velocity. The solid lines show the turnover of these systems for the forward (green) and backward (purple) reactions. For the forward reaction (MgATP + AMP 🡪 ADP + MgADP), AK's turnover was assessed using the system phosphokinase/ lactate dehydrogenase. We monitored the activity of this system in the absence of AK at different urea concentrations. The reaction was started by the injection of 2 mM ADP. Similarly, for the backward reaction (MgADP + ADP 🡪 AMP + MgATP), the relative turnover of the system hexokinase/ glucose-6-phosphate dehydrogenase was monitored after the addition of 2 mM ATP. In both cases, the addition of urea reduced turnover. However, the enzymatic velocity of the coupled system remained about three times faster than the maximum turnover of AK (dashed lines), except for 1 M urea (forward direction), where the velocity decreased strongly.

**Table S1: Michaelis-Menten parameters for the backward reaction**

| **Urea concentration [M]** | **v_max_ [s^-1^]** | **K_M_ [µM]** |
| --- | --- | --- |
| **0** | 582(±17) | 1336(±119) |
| **0.2** | 598(±26) | 1335(±176) |
| **0.4** | 604(±31) | 1435(±218) |
| **0.6** | 617(±29) | 1516(±204) |
| **0.8** | 596(±33) | 1405(±232) |

**Table S2: Change in non-linearity of steady-state time course due to alleviation of product inhibition^a^**

| **ADP concentration**  **[µM] ^b^** | **η×10^3^** | | $\frac{\boldsymbol{\eta}\mathbf{(}\mathbf{0}\mathbf{.}\mathbf{8} \mathbf{M} \mathbf{urea}\mathbf{)}}{\boldsymbol{\eta}\mathbf{(}\mathbf{0} \mathbf{M} \mathbf{urea}\mathbf{)}}$ |
| --- | --- | --- | --- |
|  | **η(0 M urea)** | **η(0.8 M urea)** |  |
| **750** | 3.92(±0.04) | 3.59(±0.13) | 1.82(±0.10) |
| **1000** | 4.36(±0.15) | 3.26(±0.05) | 0.91(±0.25) |
| **2000** | 4.72(±0.76) | 3.37(±0.16) | 0.75(±0.12) |
| **3000** | 5.14(±0.97) | 3.94(±0.14) | 0.76(±0.15) |
| **4000** | 6.73(±0.62) | 4.69(±0.68) | 0.70(±0.12) |
| **5000** | 7.89(±0.58) | 4.90(±0.27) | 0.62(±0.06) |

**^a^** Fit according to De La Cruz *et al..*^24^
**^b^** For low initial substrate concentrations (≤500 µM), where turnover numbers are small and little AMP is formed, a more reliable fit of turnover was obtained when η was set to 0.

**Table S3: Comparison of transition rates obtained by H^2^MM vs. dwell-time analysis (DTA)^a^**

| **AMP [μM]** | **Urea [M]** | ***k*_c_ [10^3^ s^-1^]** | | ***k*_o_ [10^3^ s^-1^]** | |
| --- | --- | --- | --- | --- | --- |
|  |  | **H^2^MM** | **DTA** | **H^2^MM** | **DTA** |
| **apo** | **0** | 0.5 (±0.2) | 2.1 ^b^ (±0.5) | 5.8 (±2.3) | 5.2 (±1.7) |
|  | **0.2** | 0.8 (±0.3) | 2.5 ^b^ (±0.7) | 7.3 (±3.5) | 6.7 (±0.2) |
|  | **0.4** | 0.7 (±0.5) | 2.7 ^b^ (±0.4) | 5.2 (±2.4) | 7.7 (±2.9) |
|  | **0.6** | 0.7 (±0.2) | 1.6 ^b^ (±0.1) | 4.6 (±1.3) | 4.0 (±1.3) |
|  | **0.8** | 0.6 (±0.1) | 1.7 ^b^ (±0.6) | 3.9 (±0.4) | 4.7 (±0.7) |
| **400** | **0** | 32.7 (±1.4) | 31.0 (±1.8) | 21.9 (±1.4) | 20.1 (±1.1) |
|  | **0.2** | 27.6 (±2.2) | 26.0 (±2.0) | 21.7 (±1.2) | 19.8 (±1.0) |
|  | **0.4** | 28.0 (±0.2) | 26.2 (±0.1) | 24.0 (±0.6) | 22.0 (±0.6) |
|  | **0.6** | 25.6 (±3.9) | 23.6 (±3.7) | 22.9 (±2.9) | 21.3 (±2.2) |
|  | **0.8** | 24.6 (±2.3) | 22.6 (±2.1) | 25.6 (±0.3) | 23.6 (±0.1) |
| **5000** | **0** | 46.5 (±0.4) | 42.2 (±1.7) | 28.3 (±0.9) | 24.7 (±0.6) |
|  | **0.2** | 51.1 (±1.5) | 47.5 (±1.7) | 34.6 (±1.7) | 29.9 (±1.6) |
|  | **0.4** | 46.7 (±0.3) | 41.7 (±0.1) | 33.9 (±2.2) | 30.9 (±2.4) |
|  | **0.6** | 44.4 (±0.4) | 40.5 (±1.1) | 36.7 (±1.5) | 32.6 (±2.1) |
|  | **0.8** | 40.2 (±4.0) | 36.2 (±3.1) | 36.2 (±2.4) | 32.3 (±1.3) |

^a^ error bars indicate the standard error of the mean of at least two measurements
^b^ dwell times extend the length of individual bursts (~250 μs given the chosen burst selection criteria), preventing a reliable analysis

**Table S4: Substrate concentrations used in smFRET experiments**

| ***c*_ATP_ fixed at 1 mM** | |
| --- | --- |
| **AMP [µM]** | **ADP [µM]** |
| **1** | 6.2 |
| **2.5** | 9.9 |
| **5** | 13.9 |
| **10** | 19.7 |
| **25** | 31 |
| **50** | 44 |
| **100** | 62 |
| **250** | 95 |
| **400** | 123 |
| **500** | 137 |
| **750** | 150 |
| **1000** | 160 |
| **2000** | 230 |
| **3000** | 327 |
| **5000** | 417 |
| **10000** | 576 |

**Table S5: Protein dynamics parameters used in the fitting of enzymatic velocity**

| **rate constants [ms^-1^]** | | ***k*_c_** | | | ***k*_o_** | | |
| --- | --- | --- | --- | --- | --- | --- | --- |
|  |  | **0M urea** | **0.4M urea** | **0.8M urea** | **0M urea** | **0.4M urea** | **0.8M urea** |
| **WT** | **E** | 0.5 | 0.7 | 0.6 | 5.8 | 5.2 | 3.9 |
|  | **EM** | 2.6 | 1.5 | 1.3 | 13 | 7.0 | 6.3 |
|  | **ET / ETM / ETM_i_** ^a^ | 27 | 21 | 15 | 24 | 19 | 17 |
|  | **EMT_i_** | 50 | 47 | 45 | 24 | 34 | 36 |
| **F86W** | **E** | 0.34 | 0.25 | 0.20 | 3.9 | 2.0 | 1.6 |
|  | **EM** | 0.36 | 0.24 | 0.23 | 3.6 | 2.5 | 1.5 |
|  | **ET / ETM / ETM_i_** ^a,b^ | 20 | 15 | 6.4 | 18 | 16 | 13 |
|  | **EMT_i_** ^b^ | 17 | 12 | 8.6 | 16 | 18 | 17 |
| **L107I** | **E** | 2.2 | 1.5 | 0.8 | 17 | 11 | 7.5 |
|  | **EM** | 7.5 | 5.0 | 2.6 | 38 | 34 | 18 |
|  | **ET / ETM / ETM_i_** ^a^ | 30 | 28 | 20 | 25 | 26 | 24 |
|  | **EMT_i_** | 49 | 44 | 55 | 25 | 41 | 44 |

^a^ The single-molecule experiments cannot distinguish between ET, ETM and ETM­_i_.
^b^ Differences between ET/ETM/ ETM­_i_ and EMT­_i_ in F86W are not statistically significant.

**Table S6: Parameters for the unfolding of AK measured by CD spectroscopy**

|  | ${\Delta G}_{\mathrm{fold}}^{0}$ [kJ mol^-1^] | $m$ [kJ mol^-1^ M^-1^] | midpoint [M] |
| --- | --- | --- | --- |
| without substrates | 18.8 | 6.7 | 2.80 |
| with substrates | 27.0 | 8.6 | 3.12 |
